# Optimal parameters for measuring multiband auditory brainstem responses to continuous speech

**DOI:** 10.64898/2025.12.24.696406

**Authors:** Melissa J. Polonenko, Benjamin R. Eisenreich

**Affiliations:** Department of Speech-Language-Hearing Sciences, University of Minnesota; Center for Applied and Translational Sensory Science, University of Minnesota

## Abstract

Accurate clinical hearing assessment depends on efficient, engaging measures designed to evaluate ecologically relevant stimuli. Often brief tones or narrowband noise stimuli are used, providing a useful but limited snapshot of hearing function. Including dynamic speech offers a means to capture how the hearing system encodes complex sounds critical for everyday communication. Here we describe the optimal parameters for using audiobook continuous speech with the multiband peaky speech paradigm to measure frequency-specific auditory brainstem responses (ABRs) to standard audiological octave bands from 500–8000 Hz in each ear simultaneously. Using computational modeling and direct human ABR testing in adults with normal hearing, we demonstrate that continuous speech signals with a chirp phase profile and fundamental frequency (f0) lowered to the range of 90–110 Hz evoke the largest ABR wave V amplitudes. This amplitude boost occurs when any narrator’s f0 is lowered to this optimal range, but the largest responses occur for narrators with original f0s below 170 Hz. We also confirmed that different narrator speech stimuli with these optimized parameters can evoke similarly sized ABRs, but some minor differences remain for testing time. Ultimately, optimizing phase-f0 parameters substantially sped up the median testing time to obtain robust audiobook-based multiband ABRs to within 14 minutes, thereby making this paradigm more feasible for future research and clinical translation.

## Introduction

Multiple frequency-specific auditory brainstem response (ABR) testing is an important diagnostic tool to characterize hearing loss, particularly for individuals who cannot tell us what they hear. Typically, multiband ABRs use tone pips or narrowband chirp stimuli to predict behavioral thresholds (McCreery et al., 2015; Polonenko & Maddox, 2025b; Stapells & Oates, 1997). But brief sounds are not engaging for awake individuals, and more importantly, they do not reflect all the strikingly dynamic components of continuous speech—components that are critical for speech understanding and are impacted by hearing loss. Using continuous speech stimuli may facilitate awake testing and assessment of speech encoding in new ways. The goal of this paper is to further develop one such tool – multiband ABRs to audiobooks – by optimizing its parameters to reduce test time and make it a feasible test for future research studies and clinical translation.

The recently developed multiband peaky speech paradigm measures frequency-specific ABRs to continuous speech (Polonenko & Maddox, 2021). This method simultaneously evokes ABRs for the same octave-band frequencies as diagnostic ABRs and hearing aid fittings (500 to 8000 Hz) but uses narrated stories at supra-threshold levels instead of repeated brief stimuli. These stories may engage awake individuals for longer periods of testing, making multiband peaky speech a promising method for addressing some challenges to objective testing.

But the multiband peaky speech paradigm needs to be optimized. Responses to continuous speech are very small and testing time depends on response size. Previous work showed that at least 60 minutes was required to obtain responses with a decent signal-to-noise ratio (SNR) for all 10 bands (2 ears x 5 octave bands) for most adults with normal hearing (Polonenko & Maddox, 2021). Some adults require less time but when creating an assessment paradigm we need to know what acquisition times to expect for most people. Reducing the testing time is necessary for feasible use in research, with different populations, and for eventual translation to the clinic. Our major goal of this work was to optimize parameters to increase response size to address this time limitation. Previous work shows that responses are smallest for the low frequency octave bands and depend on the narrator (Polonenko & Maddox, 2021). Thus, two important parameters can be modified to increase response size and thus speed up testing time: 1) phase profile; and 2) narrator fundamental frequency (f0).

The first parameter we modified is the phase profile. Peaky speech was originally created by aligning the phase of all speech harmonics to zero (cosines) at each period of the f0, similar to the phase profile of click and tone pip stimuli of traditional ABRs. The zero-phase profile gave canonical responses that characteristically became broader, smaller, and later for the lower frequency bands, corresponding to the poorer synchrony and longer traveling wave delays for the cochlear apex compared to the base. Chirp-phase profiles compensate for the cochlear delays by introducing frequency-dependent phase delays, which leads to more synchronization along the cochlea and larger ABRs than to clicks or tone pips in adults (e.g., Dau et al., 2000; Elberling & Don, 2008), as well as in infants and children with normal hearing and hearing loss (Cobb & Stuart, 2016; Maloff & Hood, 2014; Zirn et al., 2014). Chirp-phases are easily integrated into multiband peaky speech by applying an all-pass phase function to the stimuli. There are different chirp-profiles and we chose to use the level-independent CE-chirp (Elberling et al., 2007) because the connection between the published levels for level-dependent chirps and those of our speech bands is not obvious.

The second parameter that could improve response size and speed up testing time is the narrator f0. The f0 is akin to stimulus rate, which is a well-known parameter for the brief clicks and tone pips used in traditional ABRs. Slower stimulus rates give larger ABRs due to less neural adaptation and more time for all neurons to have a refractory period and synchronously fire together (e.g., Burkard et al., 1990; Burkard & Hecox, 1983; Don et al., 1977; Jiang et al., 2009; Polonenko & Maddox, 2022). In the first proof-of-concept study of peaky speech (Polonenko & Maddox, 2021), we showed the male narrator evoked larger responses with better SNR than the female narrator, and then established that this effect was driven by f0―rather than gender―in a follow-up study that systematically changed the f0 of the same two narrators for broadband ABRs (Polonenko & Maddox, 2024). However, the optimal f0 range is unknown and needs to be evaluated for multiband ABRs to continuous speech. Furthermore, not all labs and clinics will have access to a narrator’s voice with a very low f0; thus, we also aimed to determine if we could lower any narrator’s f0 to boost response size and therefore shorten testing time, as well as allow for a wider audiobook selection.

In this paper we systematically examined the optimal f0-phase combination for evoking the largest―and thus fastest to record―multiband peaky speech ABRs to make this paradigm more practical. We first used computational modeling to sample a broad subspace of f0-phase combinations with multiple audiobooks with different narrators to narrow down the most important conditions to verify in the human electroencephalography (EEG) experiments. Computational modeling gives a decent approximation to the overall amplitude patterns in ABRs to different stimuli (e.g., Dau, 2003; Polonenko & Maddox, 2024; Stoll & Maddox, 2023; Temboury-Gutierrez et al., 2024) and has shown ABR changes with f0 changes to broadband peaky speech (Polonenko & Maddox, 2024). In both the modeling and EEG studies we found that chirp-phase boosts response size of the low frequency bands and lower f0s provide a more global boost irrespective of the original f0; however, the resulting amplitudes were largest if the original f0 was <170 Hz and lowered to an optimal f0 ∼90-110 Hz. The chirp-boost alone cut the median testing time in half to acquire all responses with decent SNR, and the additional f0 boost reduced test time even further to so most participants required under 20 minutes. We finally verified that different narrators could be used with the same optimized f0-chirp parameters, but some small differences remain in ABR size and test time. Overall, optimizing parameters improved response size and necessary testing times for multiband peaky speech ABRs, making this paradigm more useful as a potential tool to assess speech encoding.

## Methods

### Stimuli

#### Audiobook selection

Audiobook recordings in the public domain were selected from the children’s fiction genre of the LibriVox Collection (https://librivox.org/). Audio recordings were chosen based upon story content and narrator voice to cover a wide range of original f0s. Selected stories were screened to ensure there was one single narrator for the entirety of the story, a minimum recording sampling rate of 44.1 kHz, and a minimum bit depth of 64 kb. The final set of audio stimuli consisted of twelve stories with average narrator fundamental frequencies ranging from 113 to 215 Hz: *Alice’s Adventures in Wonderland (version 3)* (Yearsley, 2007), *The Wonderful Wizard of Oz* (Hall, 2007), *Doctor Dolittle’s Zoo* (Chenevert, 2022), *The Time Machine* (version 3) (Nelson, 2011), *Alice’s Adventures in Wonderland* (version 7) (Franklin, 2020a), *Through the Looking Glass (version 6)* (Franklin, 2020b), *The Adventures of Peter Cottontail (version 2)* (Somers, 2017), *Toto’s Merry Winter* (Somes, 2015), *The Clock Strikes Thirteen* (Adam, 2015), *The Adventures of Maya the Bee* (Bush, 2009), *Billy Whisker the Autobiography of a Goat* (Hester, 2011), *The Sea Fairies* (Bieber, 2008).

#### Pre-processing and frequency shifting

All audio was preprocessed using custom Python scripts using the Parselmouth (Jadoul et al., 2018) implementation of Praat (Boersma & Weenink, 2018). Preprocessing consisted of limiting periods of silence to under 0.5 s, resampling to a rate of 48 kHz, and slicing the audio into 10 s segments. Spectrograms of the audio segments were examined to confirm absence of filter combing or abnormalities within the frequency range of 20 Hz–20 kHz.

To explore how changes to f0 impact ABR waveforms, we used the original f0 and then shifted the f0 of each of the 12 stories. For the computational modeling, 15 minutes of audio was used. The mean, minimum, and maximum f0 were extracted for each 10 s audio segment. The extracted mean f0 was then compared to the desired target frequency to determine the semitone difference using the equation: 12*log*_2_(*original f*0⁄*shifted f*0). Using the sound manipulation class within Praat (Boersma & Weenink, 2018), the original audio pitch tier information and desired target frequency were resynthesized using the overlap method to either raise or lower the pitch of the audio signal to the target frequency. In additional to the original f0, each narrator had their f0 shifted to cover the range from 70–210 Hz in 10 Hz steps. For the EEG experiments, ∼60 minutes of each narrator was processed and shifted down to the optimal f0 (details about optimal f0 are described below).

#### Creating peaky speech and phase profiles

Both the original f0 and shifted f0 audio segments were re-synthesized into multiband peaky speech. Both zero-phase and chirp-phase stimuli were created in the same script and could use the same regressors in the deconvolution to derive the multiband ABRs. Full details can be found in Polonenko & Maddox (Polonenko & Maddox, 2021), but a description is given below.

There are two main design goals of multiband peaky speech. First, while preserving the speech’s defining spectrotemporal features we create sharp peaks in the pressure waveform to maximally drive the ABR with impulse-like stimuli. Supplemental Figure 1 compares unaltered and re-synthesized stimuli with zero-phase and chirp-phase. Second, we create speech comprised of independent frequency bands to yield separate ABR responses from the same audio and recording.

During voiced speech the vocal folds rapidly open and close, and the timing of these glottal pulses sets the dynamic f0 of speech. Frequencies at integer multiples of f0 (harmonics) also vary in amplitude and phase over time. To make speech “peaky” we align the phases of the harmonics so that they all peak at each glottal pulse, creating one large peak when they are summed together. This is the zero-phase profile. To make speech with multiple bands, we create sets of harmonics based on a randomly modified f0 for each octave-band which makes the bands independent over time. To do this, we extract the glottal pulses and then identify the voiced sections based on an inter-pulse interval <17 ms, corresponding to an f0 >60 Hz. To re-synthesize the f0 waveform, *f*_0_(*t*), a phase function, *φ*(*t*), is created that increases smoothly by 2*π* between the glottal pulses. Based on random frequency shifts, *f*_Δ_, of up to ±1 Hz over the duration of the stimulus, as well as a random starting phase, *θ*_Δ_, we modify this phase function to make it independent according to the formula: 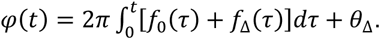 We calculate the speech’s spectrogram to determine the amplitudes of each frequency at each time point, *A*(*t*, *f*), and then create the new f0 waveform as *h*_0_(*t*) = *A*[*t*, *f*_0_(*t*)] cos[*φ*(*t*)] and each *k*^th^ harmonic at frequency (*k* + 1)*f*_0_ as *h*_*k*_(*t*) = *A*[*t*, (*k* + 1)*f*_0_(*t*)] cos[(*k* + 1)*φ*(*t*)], which are summed to create the re-synthesized voiced speech. This process is repeated for five octave-bands (center frequencies of 0.5, 1, 2, 4, 8 kHz) for each ear. Each of the 10 independent sets of voiced speech is band-pass filtered to their respective octave-bands and then the bands for each ear are mixed together. For chirp-phase, the re-synthesized voiced speech is then all-pass filtered by convolving with a chirp-phase profile. We used the level-independent CE-Chirp (Elberling et al., 2007; Elberling & Don, 2008) with this formula: *τ* = *kf*^−*d*^ where *τ* is the delay in seconds and *f* is the frequency in Hz and *k* and *d* are constants equal to 0.092 and 0.4356 respectively. Finally, the re-synthesized voiced speech is cross-faded back and forth with the original unvoiced speech and filtered up to 11.36 kHz, above which we used the unaltered speech to improve the quality of voiced consonants.

### Computational modeling

Modeled ABRs for 10 bands (0.5, 1, 2, 4, 8 kHz bands each in 2 ears) were generated from our multiband peaky speech stimuli using custom written python scripts and well-known computational models of the auditory nerve and periphery. To fully explore the entire parameter space, 15 minutes (90 segments) of audio for each phase × f0 × narrator combination was used. As done before, the stimuli were up-sampled to 100 kHz and passed into the python implementation of the peripheral model (Rudnicki et al., 2015; Zilany et al., 2014) with a standard flat audiogram of 0 dB HL and high spontaneous firing rate auditory nerve fibers with characteristic frequencies across ⅙-octave intervals from 0.125–16 kHz (Kulasingham et al., 2024; Polonenko & Maddox, 2024; Shan et al., 2024). The auditory nerve fiber responses from the cochlear model were then input into the brainstem model (Verhulst et al., 2018) to give the simulated EEG. Modeled ABRs were then derived through deconvolution in the same way as the human EEG, as described below in the analysis section. Wave V amplitudes and latencies were calculated using automated peak peaking and were verified by human researchers (see below for details).

### Human EEG experiments

Next we verified the chirp-phase and f0 effects identified by the computation modeling and confirmed the estimated needed recording times in a series of three human EEG experiments: 1) zero- versus chirp-phase stimuli of a narrator with a low original f0; 2) chirp-phase stimuli made from narrators with a low and high original f0 that were also shifted down to an optimal f0 of 100 Hz; and 3) chirp-phase stimuli made from three narrators with original f0s <170 Hz that were shifted down to 90 Hz to verify different stories with optimized parameters could evoke similarly sized multiband ABRs, thus allowing a variety of audiobooks to be flexibly used.

#### Participants

All participants gave written informed consent before participating and underwent a protocol approved by the University of Minnesota Institutional Review Board (Study #17008). A different group of adults were recruited for each of the 3 experiments, although 8 participants completed more than one experiment. A hearing screening confirmed hearing thresholds ≤20 dB HL from 0.25–8 kHz in both ears.

A total of 62 participants aged 18–57 years were recruited, although data from 7 participants were excluded due to experimental issues resulting in no responses (experimenter forgot to hit “record” for the last hour of recording; the earphone was plugged with earwax in one ear or fell out midway through the recording) or very noisy responses (n=3). Data was deemed noisy based on replications. Replications comprised of half the data split by odd and even trials were derived and compared using Spearman correlation coefficients over a 9 ms window (for details see the section on optimal analysis parameters below). Data was deemed too noisy and excluded if the median correlation coefficient across the 10 bands was less than 0.4. With this criterion, data from 3 participants in experiment 2 were excluded. We confirmed that the ABRs had poor morphology and experiment notes from the recording session indicated abnormally noisy EEG despite good impedances and efforts by the experimenter to reduce noise. These participants came to the lab during winter months with a lot of static and dry air, and the booth was often cold. They were provided with a blanket, but the static and muscle tension may have contributed to the noisy EEG. We have since installed larger humidifiers in the lab to control for static during cold winter months.

The final numbers for experiments 1–3 were 15, 25, and 15 respectively for a total of 55 participants. More participants were recruited for experiment 2 to ensure sufficient comparisons could be made due to the expected smaller ABRs evoked by the narrator with the high original f0. The mean ± SD age for each group was 21.6 ± 5.6 years for experiment 1, 21.8 ± 5.9 years for experiment 2, and 20.3 ± 2.0 years for experiment 3.

#### Stimuli for EEG experiments

The total minutes per condition varied across the experiments based on the results of these and previous experiments and to keep a recording session under 2.5 hours. Ideally, responses from normal hearing adults could be obtained within 35 minutes (the shortest recording time per condition).

For experiment 1, 60 minutes each of zero- and chirp-phase stimuli were recorded (total 2 hours) based on our previous work with multiband ABRs (Polonenko & Maddox, 2021). We chose a narrator with a lower original f0 of 124 Hz (Hall, 2007) to ensure larger ABRs were recorded, especially with the zero-phase stimuli.

For experiment 2, two narrators were chosen: the same narrator with an 124 Hz f0 as in experiment 1 (Hall, 2007) and another narrator with a higher 180 Hz f0 (Somes, 2015). Conditions included these two original f0s and each narrator with an f0 shifted down to 100 Hz, determined as optimal based on the modeled ABRs. Based on the chirp-phase results from experiment 1, 35 minutes each of these four conditions were recorded, for a total of 2 hours and 20 minutes.

For experiment 3, three narrators were chosen with f0s <170 Hz, one of which was the same narrator that was used in experiments 1 and 2. The original f0s were 124 Hz (Hall, 2007), 130 Hz (Franklin, 2020a), and 147 Hz (Chenevert, 2022). 35 minutes of each narrator were recorded for a total of 105 minutes.

#### ABR recording

Participants sat in a reclined chair in a darkened sound booth while listening to the audiobooks and wearing an EEG cap. Stimulus conditions and 10-s audio segments were randomized and played via custom python scripts using expyfun (Larson et al., 2014). The multiband peaky speech stimuli were presented at a sampling rate of 48 kHz through an RME digiface USB soundcard coupled to ER-2 insert earphones. The stimuli were calibrated relative to a 1000 Hz pure tone (Polonenko & Maddox, 2021) and confirmed to have an overall root-mean-square (RMS) level between 65–70 dB SPL through a sound level meter coupled to a 2cc coupler. The participant or experimenter could pause and resume the experiment if needed.

ABRs were recorded with 4 passive Ag-Cl electrodes placed at Fz (active, non-inverting), A1 and A2 (reference, inverting), and Fpz (ground). Impedances were kept below 3 kohms but most were 1 kohm. Continuous EEG was recorded at a sampling rate of 10 kHz using a BrainVision actiCHamp+ amplifier and two differential EP pre-amplifiers connected to BrainVision Recorder software. Triggers marking the onset (1) and offset (2) of each 10-s audio segment, along with triggers indicating the trial information (binary trigger converted to 4’s and 8’s), were sent via the soundcard to a custom trigger box (Maddox, 2020) that converted the triggers to a TTL signal that was then sent to the EEG amplifier to synchronize the timing of stimuli and EEG with sub- µs precision.

### Analyses

#### ABR pre-processing and derivation

Modeled and measured ABRs were derived using custom python scripts that incorporated the *mne* package (Gramfort et al., 2013) as described in Polonenko & Maddox (Polonenko & Maddox, 2021). Modeled and measured EEG were bandpass filtered from 30–2000 Hz using a 1st order causal Butterworth filter and notch filtered at odd multiples of 60 Hz (causal infinite impulse response, IIR) to remove electrical line noise. The data were epoched from 500 ms before the stimulus onset (marked by a trigger) to 500 ms after the 10-s trial for a total of 11 s per epoch. Another trigger marked the stimulus offset, which was used to correct the regressor for subtle clock drifts between the soundcard and the EEG amplifier over the 10-s trial. This was done during resampling of the glottal pulse train from 48 kHz to match the EEG sampling rate of 10 kHz. The regressors were created by placing 1’s (impulses) at indices corresponding to the resampled timing of the glottal pulses of the peaky speech, then zero-padding the first and last 500 ms to match the EEG epoch. Because of the random and independent timing used to create the different bands of the peaky speech, the same EEG could be used with each band’s pulse train regressor to derive waveforms to each of the 10 bands (5 bands in 2 ears).

The ABR for each band was derived by deconvolving the EEG with its respective pulse train, done via frequency domain division for efficiency (Polonenko & Maddox, 2021). The numerator was the cross-spectral density (cross-correlation in the time domain) of the pulse train (*x*) and EEG (*y*), and the denominator was the power spectral density of the pulse train (autocorrelation in the time domain), according to the formula: *w* = *F*^−1^{〈*F*{*x*}^∗^*F*{*y*}〉⁄〈*F*{*x*}^∗^*F*{*x*}〉}, where ⟨⟩ indicates average across trials, * represents complex conjugation, *F* the fast Fourier transform and *F*^−1^ its inverse. As effectively done in prior studies (Polonenko & Maddox, 2019, 2021, 2022, 2024) to improve signal quality each epoch was weighted by the inverse of its variance relative to the summed variance of all epochs to reduce noise and avoid excluding epochs based on noise criteria. Due to the circular nature of the FFTs, the last 500 ms and first 500 ms were concatenated together to give the ABRs from [-500, 500] ms with 0 ms marking stimulus onset. The middle 10 s was discarded. This was repeated for each of the 10 bands. We also calculated and then subtracted out a common component response using an average of responses to 5 sham pulse trains that are not presented as stimuli. This allows us to extract truly frequency-specific responses by subtracting coherent activity resulting from pulse trains of each band starting and stopping together during voiced segments (Polonenko & Maddox, 2021).

#### ABR wave V metrics

Modeled and measured ABR wave V peaks were automatically picked and then visually verified by the authors. In some cases, minor adjustments were made to the auto-pick parameters to ensure wave V was selected rather than an earlier component wave, especially for the modeled responses that showed more distinct earlier component waves I and III. Auto-picks were done to allow for comparing multiple filtering parameters (see section below). The starting parameters for the auto-picks were determined from the grand average ABRs for each experimental condition by finding the maximum amplitude (wave V) and minimum amplitude (wave Vp, or trough of wave V) from 2 to 20 ms. Then individual narrator (modeled) or participant (measured) ABR wave V metrics were calculated taking the grand average wave V and Vp peak latencies with a range from -1 to +2 ms and finding the respective amplitude maximum and minimum across these latency ranges. Every auto-pick was visually confirmed. For a few participants the range was adjusted to -0.25 to +2 ms to avoid auto-picking wave III instead of wave V. Wave V amplitude was calculated as the difference in amplitude between wave V and Vp (i.e., the peak-to-trough amplitude). All waveforms with auto-peaks are provided in Supplemental Figures 3-17 (modeled ABRs: Supplemental Figures 3–8; measured ABRs: Supplemental Figures 9–17).

Manually picked wave V metrics were also done for the 150–1500 Hz bandpass filtered measured data, as a starting place based on previous work (Polonenko & Maddox, 2021). Three inexperienced students and the experienced first author picked peaks. An intraclass correlation coefficient type 3 analysis indicated that these manually picked peaks showed good agreement with each other and with the auto-picks for these filter settings (all *ICC3* >0.981 for peak amplitude and >0.831 for peak latency).

#### SNR and test time metrics

The main aim of this work was to increase wave V amplitudes to obtain better response SNRs so that less time is required to record a robust response. Testing time was estimated as the time to reach 0 dB SNR, as done before (Polonenko & Maddox, 2019, 2021, 2022, 2025a). To do this, the ABR SNR was calculated as the response variation, 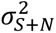 over a 9 ms window encompassing the ABR response (see section below) relative to the average variation across similarly sized windows from -480 to -20 ms of the pre-stimulus baseline, 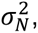 according to the following equation: 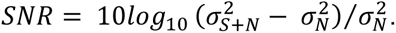 Then the SNR for 1 minute of recording was estimated by: 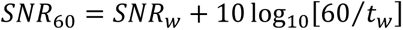 where *t*_*w*_ is the duration of the recording in seconds (2,100 s for 35 minutes and 3,600 s for 60 minutes). The *SNR*_60_ was then used to calculate the time to reach 0 dB SNR for each participant, *t*_0*dB SNR*_, by 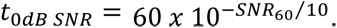 An extrapolation up to 120 minutes (twice the recording time) was taken as the maximum estimated required test time. Time was also characterized by a cumulative density function (CDF) to determine what recording time was necessary for a certain percentage of participants to have all 10 waveforms reach the 0 dB SNR criterion. Ideally, 100% of participants would achieve this stopping criterion in under 30 minutes. The time noted on the CDF would be for the slowest waveform of the 10 bands to achieve 0 dB SNR, representing the point in time an experimenter could stop the testing.

SNR and test time were not calculated for the modeled ABRs because there was very low noise in the simulated EEG and pre-stimulus baseline. Any calculation would over-estimate the SNR and under-estimate the required test time for measured EEG. Thus, only wave V metrics are given for the modeled data.

#### Optimized post-processing parameters

Previous work suggested that 150–1500 Hz was the optimal bandpass filter parameters for multiband ABRs (Polonenko & Maddox, 2021). Thus, initially wave V metrics were evaluated with these filter parameters (including the manually picked wave V peaks). However, this paper aimed to find the optimal stimulus and analysis parameters for measuring multiband speech ABRs, and so a systematic approach was taken to determine the optimal filtering and SNR analysis window parameters. ABRs were bandpass filtered with a zero-phase 1^st^ order Butterworth filter with a low-pass cutoff of 1500 Hz and high-pass cutoffs from 30–150 Hz in 10 Hz steps. The latency range window to calculate the response SNR and time-to-0dB varied from 8.0–12.5 ms in 0.5 ms steps.

For chirp-phase stimuli the SNR analysis window started at 0 ms. For zero-phase stimuli, the ABR waveforms started at progressively later latencies with decreasing frequency (and more apical cochlear stimulation). Thus, the analysis window was shifted for each octave band to encompass the ABR response in a similar way for the zero-phase and chirp-phase responses to ensure appropriate SNR and timing comparisons between the two phases. To do this, the shifts were calculated as the difference in the grand average wave V peak latencies. This resulted in the analysis windows for 0.5, 1, 2, 4, and 8 kHz bands starting at 5.9, 4.6, 3.2, 2.5, and 1.9 ms respectively for the measured ABRs.

The optimal bandpass filter parameters and SNR analysis window combination was then determined based on the area under the CDF curves (AUC), normalized to 1 s. The closer the AUC to 1, the more participants who had all 10 waveforms at 0 dB SNR in less time. While several parameters gave comparable AUCs, the overall optimal parameters across all 3 experiments included a bandpass filter of 80–1500 Hz and a 9.0 ms SNR analysis window.

#### Optimized f0 shift

Modeled ABR wave V amplitude was used to determine the optimal f0 shift for each narrated audiobook. First, the overall optimal shifted f0 range was determined by averaging the modeled ABRs across ear, narrator and phase, and then calculating the range of f0s over which the wave V amplitude was at least 90% of the peak wave V amplitude. From the overall average, the optimal f0 range was 80–130 Hz. Then this range was used to determine the optimal f0 for each narrator. This was done by taking the mean wave V amplitudes across phase, finding the f0 with the maximum amplitude for each octave band, and then calculating the mode optimal f0 across octave bands to give a single optimal f0 for that audiobook. The optimal f0 was visually confirmed and is displayed in Supplemental Figure 2. The optimal f0 was most frequently 100 Hz (n=8).

#### ABR boosts

The increase in ABR amplitude and reduction in test time were assessed by taking the ratio of one condition to another. Thus, the boost is given in terms of “times larger” or “times faster”. The SNR boost was calculated as the dB difference between conditions.

#### Statistical Analyses

Shapiro-Wilk tests of normality confirmed that most metrics were not normally distributed. This is not surprising given that most of the statistical analyses were performed on the ratio (proportional) data. Thus, non-parametric statistics were conducted. Comparisons across octave bands or between conditions were first analyzed using the Friedman Chi-Squared test, followed by Wilcoxon Signed-Rank post-hoc tests. False discovery rate (FDR; Benjamini & Hochberg, 1995) was used to correct p-values for multiple comparisons. A proportions Z-test was done to compare whether a greater proportion than 50% of participants had faster test times (time-to-0 dB) with chirp-versus zero-phase or with shifted versus original f0.

## Results

We first determined the optimal range of parameters using computational modeling and then confirmed the main phase- and f0-effects with human EEG experiments. The main outcomes were response size and the testing time required to achieve a robust response, determined as 0 dB SNR (Polonenko & Maddox, 2019, 2021, 2022, 2025a). Ultimately, the goal was to determine a set of parameters that allows us to measure robust responses in minimal test times to create a feasibly translational assessment tool. A CDF was used to show the required test times for proportions of participants to complete an exam with all 10 waveforms (5 ABRs in 2 ears) reaching the 0 dB SNR criterion.

### Modeling suggests larger ABRs are evoked by audiobooks with narrators who have lower f0 and chirp-phase profiles

Our modeling approach focused on finding the optimal parameter space within which multiband peaky speech will produce the largest ABRs across the octave bands of 0.5–8 kHz. First, we modeled 12 audiobooks with original f0s ranging from 124 to 215 Hz using zero- and chirp-phase profiles to the multiband peaky speech. Half the narrators had f0s <170 Hz. Modeled wave V amplitudes, averaged across ears, are given in the top row of Figure 1 and show two important parametric boosts: lower f0 and chirp-phase. Across all octave bands, ABR amplitudes increased with decreasing narrator f0. Furthermore, those narrators with f0s <170 Hz had a larger chirp-phase benefit for the lowest (0.5 kHz) octave band. Chirp-phase appeared to minimally change ABR amplitude for the 1–8 kHz bands.

**Figure 1.**
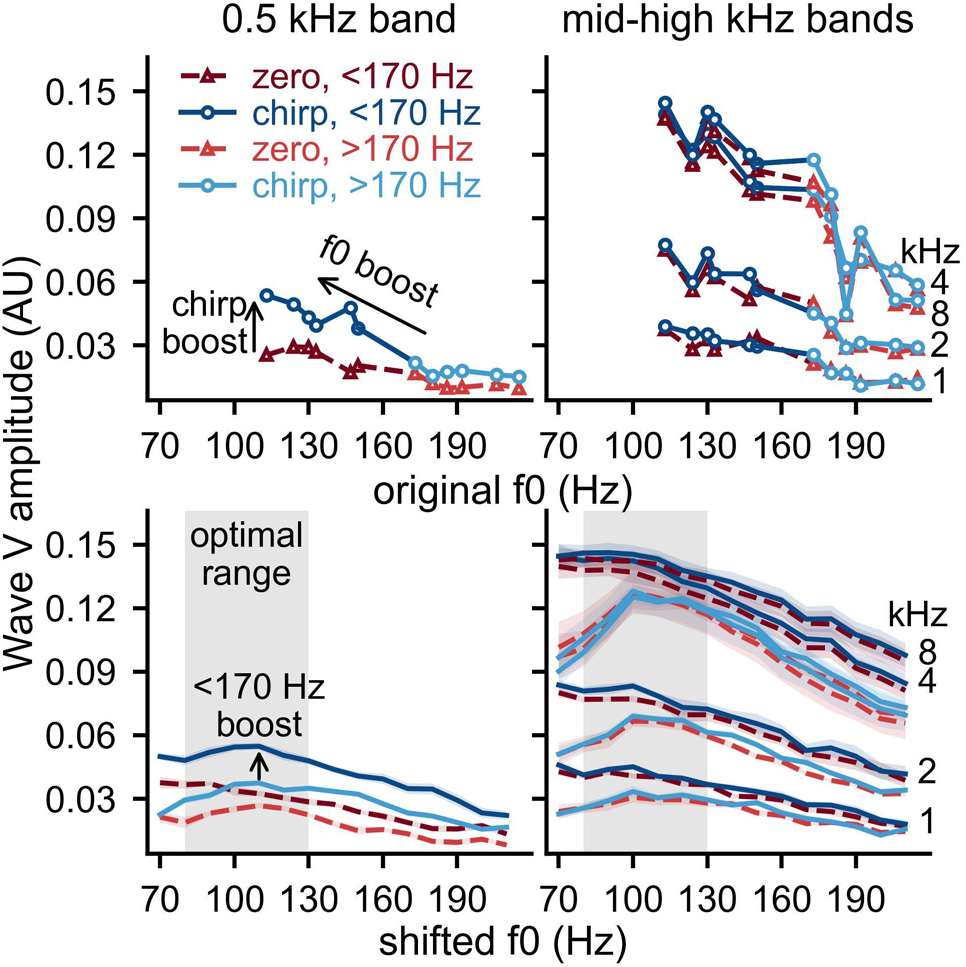
Modeled wave V amplitudes for stories with original f0s (top) and shifted f0s (bottom). Data are averaged across the stories that have original f0s <170 Hz and >170 Hz. The shaded region denotes the range used to select the optimal f0.

Next, we investigated whether any narrator’s voice could be shifted down to a lower f0 and evoke larger ABRs and confer the same chirp-phase benefit for lower shifted f0s. The bottom row of Figure 1 shows that shifting f0 lower indeed increased modeled wave V amplitudes for all narrators, but the amplitudes remained larger if the narrator had an original f0 <170 Hz (darker lines are higher than the lighter lines). This advantage of using narrators with lower f0s and further shifting the f0 lower is also seen with the chirp-phase benefit in the 0.5 kHz band.

The lowered f0 and phase boost trends were further analyzed in Figure 2. The modeled mean ABR waveforms are shown in Figure 2A and the respective auto-picked wave V peak amplitudes and latencies in Figure 2B. Data are shown for original f0s and each phase (left and middle panels of A) and the optimal f0s with chirp-phase (right panel of A). While modeled in Figure 1 and Supplemental Figure 2, data for zero-phase and optimal f0 are not shown because this set of parameters is sub-optimal. Zero-phase wave V peaks typically became earlier and larger with increased octave band. Chirp-phase wave V amplitudes also increased with higher octave bands, but the latencies were more similar across octave bands, with a subtle increase in latency for higher octave bands. Both zero- and chirp-phase wave V peaks were earlier and larger for lower shifted f0s than higher shifted f0s, consistent with the effects seen with broadband speech (Polonenko & Maddox, 2021, 2024).

**Figure 2.**
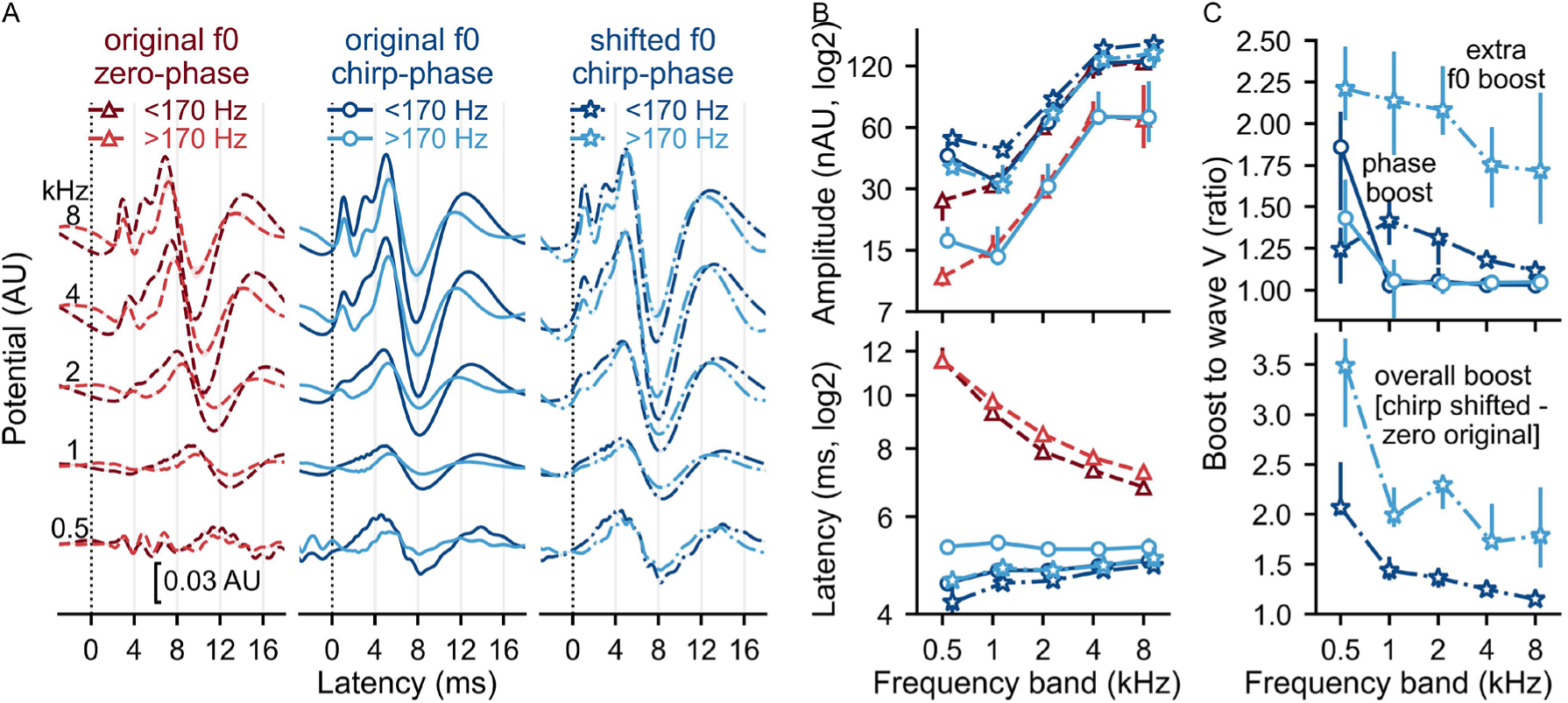
Modeled ABRs to parameter changes. (A) Waveforms for different phases and f0s and the corresponding (B) wave V peak amplitudes (top) and latencies (bottom). (C) Top plot shows boosts to wave V amplitude when changing phase (chirp – zero) and then by shifting the f0 down (optimal – original for chirp phase). Bottom plot shows the overall boost by changing both parameters (optimal shifted f0 with chirp-phase – original f0 with zero-phase).

Of greatest importance for our main goal is the extent of the boosts upon changing parameters, shown as ratios in Figure 2C. The top row shows the separate boosts for phase and f0, and the bottom row shows the combined f0-phase boost. Simply changing the underlying phase profile from zero- to chirp-phase, the least perceptually noticeable and easiest parameter change to incorporate when creating multiband peaky speech, produced a median amplitude boost to the 0.5 kHz band of 1.86× larger for f0s <170 Hz and 1.43× larger for f0s >170 Hz but minimal changes (ratios ∼ 1.03×) to the 1–8 kHz bands for all f0s. In addition to using chirp-phase, further lowering the f0 to the narrator’s optimal f0 (most commonly ∼100 Hz) gave an additional boost across all octave bands ranging from a median of 1.12× to 1.42× larger for f0s <170 Hz and 1.72× to 2.21× larger for f0s >170Hz. Taken together, changing both parameters from zero-phase and original f0 to chirp-phase and lowered optimal f0 produced an estimated median wave V amplitude boost ranging from 1.15× larger for the 8 kHz band to 2.06× larger for the 0.5 kHz band for narrators with original f0s <170 Hz, and 1.79× to 3.49× larger respectively for the 8 and 0.5 kHz bands for narrators with original f0s >170 Hz.

Although there are overall combined f0-phase boosts, we next verified each boost separately in human EEG experiments to ensure we had sufficiently robust ABRs to compare and to determine the relative time-saving benefits conferred by each successive parameter change. Because previous work with multiband peaky speech indicated at least ∼60 minutes was required for robust responses with a male voice (lower f0) and zero-phase (Polonenko & Maddox, 2021), we first conducted an experiment with zero- versus chirp-phase using a narrator with a lower original f0. Then we conducted an experiment with chirp-phase and both original and lowered f0s for two narrators, one with an f0 <170 Hz and one with an f0 >170 Hz. Finally, we ran a third experiment to evaluate whether using different narrators with f0s <170 Hz conferred similar ABR amplitude boosts and time-saving benefits, thereby allowing a varied set of audiobooks to be used with this paradigm.

### Chirp-phase boosts multiband responses and cuts the median testing time in half

First, we examined the ABR wave V amplitude and response SNR boosts, as well as the corresponding time-saving benefits to simply changing the underlying phase profile, with all data shown in Figure 3. This represents the simplest, and least perceptually different, parameter change for multiband peaky speech.

**Figure 3.**
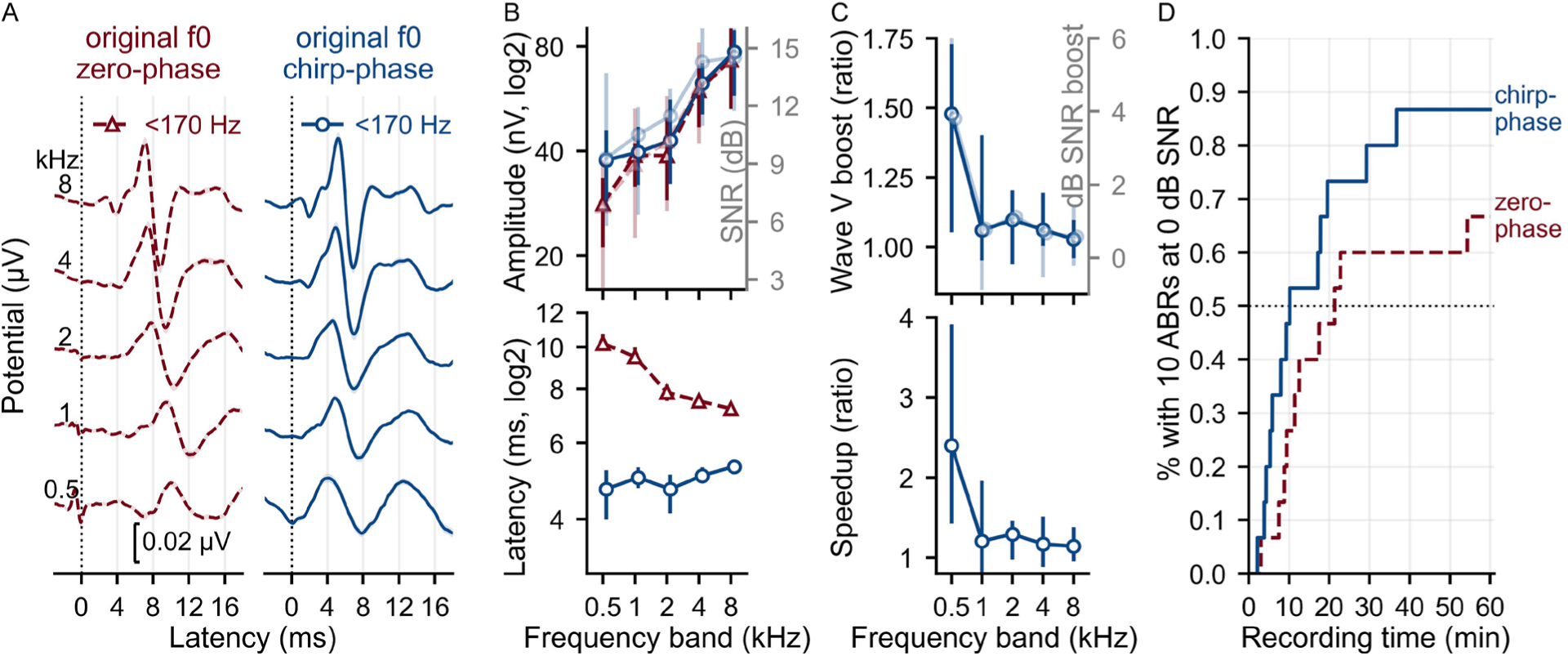
Measured ABRs to changing phase for n=15. (A) Mean ± SE (shading) waveforms for zero- and chirp- phase, along with their corresponding median (B) wave V peak amplitudes (top) and latencies(bottom). The signal-to-noise ratio (SNR) over 0-9 ms is also displayed along with the amplitude. (C) Chirp-phase boosts wave V amplitude and response SNR over 0-9 ms (top) and the associated speedup ratio of testing time calculated from the SNR (bottom). All error bars represent the 25^th^ to 75^th^ percentiles. (D) Cumulative density function of the proportion of participants who had all 10 waveforms (5 bands in 2 ears) reach 0 dB SNR across recording time in minutes. The dotted horizontal line denotes the point at which half the participants achieved this criterion.

Figure 3A and B show the mean waveforms, response SNR over a 9 ms latency window, and ABR wave V peak amplitude and latency. Wave V latencies were earlier and more similar across frequency band for chirp- then zero-phase, but they still differed across frequency bands (*χ^2^*(4)=34.73, *p*<0.001), suggesting the CE-chirp did not fully account for the cochlear traveling delays for the speech stimuli. ABR response SNRs and wave V peak amplitudes for both phases increased with increasing frequency band in a similar way (SNR: zero-phase *χ^2^*(4)=65.30, *p*<0.001, chirp-phase: *χ^2^*(4)=49.95, *p*<0.001; wave V amplitude: zero-phase *χ^2^*(4)=82.43, *p*<0.001; chirp-phase *χ^2^*(4)=75.79, *p*<0.001).

To compare responses between the phases, the ratio of wave V amplitude and the SNR difference were calculated, as well as the speedup ratio of time to reach 0 dB SNR criterion across frequency bands (Figure 3C). Using chirp-phase boosted wave V amplitude by a median of 1.48× larger (inter-quartile range, IQR: 1.05–1.73×) for the 0.5 kHz frequency band, which was significantly greater than 1 (Wilcoxon Signed-Rank test, *Z*=418, *p*<0.001) and larger than the median boosts of 1.03–1.10× larger for the other frequencies (*χ^2^*(4)=16.72, *p*=0.002; *Ζ*≤92, *p*≤0.007). This corresponded to a median SNR boost of 3.81 dB (IQR: 1.53–6.01 dB, *Z*=397, *p*=0.001) and translated into a speedup ratio of 2.40× faster (IQR: 1.42–3.91×; *Z*=435, *p*<0.001) for the 0.5 kHz band. In fact, 83.3% of ears had faster times to 0 dB SNR for chirp- than zero-phase stimuli for the 0.5 kHz band, which was significantly greater than 50% (*Z*=4.1, *p*<0.001). Even though the chirp-phase boosts (to wave V, SNR and test time) were greatest for the 0.5 kHz frequency band, the relatively smaller boosts for the other frequencies resulted in faster times for at least half the ears (63.3–70%; *Z*≤2.39, *p*≥0.042).

While frequency octave band-based boosts showed the specific phase effects, the most important outcome is their contribution to an efficient complete exam (10 ABRs at ≥ 0 dB SNR: 5 ABRs in 2 ears). As a result of the chirp boosts, participants achieved the 0 dB SNR criterion for all 10 waveforms a median of 2.24× faster (IQR: 1.22–3.02×), corresponding to a median of 10.2 minutes (IQR: 5.6–24.4 minutes) for chirp-phase compared to 21.3 minutes (IQR: 10.4– 120.0 minutes) for zero-phase, which can be seen as the CDF crossing the dotted black line in Figure 3D. In addition to cutting the median test time in half, the chirp benefits became more pronounced as testing time increased, reflected by divergence of the two CDFs after the median test times. The percentage of participants meeting the criterion for chirp- versus zero-phase was 73%:50% after 21 minutes, 80%:60% after 30 minutes, and finally 87%:67% after a maximum recording time of 60 minutes. Thus, most participants could be tested in under 20–30 minutes with chirp- but not zero-phase stimuli.

### Lowering f0 provides additional ABR boosts and time saved

Next, we verified the effect of lowering f0s for narrators with original f0s below/above 170 Hz on response SNR and wave V amplitudes, and evaluated the corresponding test times for a complete assessment (10 waveforms at 0 dB SNR). The two narrators had original f0s of 124 Hz and 180 Hz, and an optimal lowered f0 of 100 Hz. To keep the overall recording session under 2.5 hours, we recorded 35 minutes each of the four conditions (original and optimal f0s × f0s <170 and >170 Hz), all of which had chirp-phase profiles.

ABRs across all frequency bands were clearly larger and peaked earlier for the narrator with a lower original f0 but became more similar when the f0 was lowered for both narrators (waveforms in Figure 4A and wave V peak amplitude and latency and response SNR in Figure 4B). Consequently, the wave V amplitude, SNR, and speedup ratios were larger for the >170 Hz narrator than the <170 Hz narrator (Figure 4C). Furthermore, the boosts were much greater for the low-frequency bands for the >170 Hz narrator (*χ^2^*(4)=86.53, *p*<0.001; *Ζ*≤315, *p*≤0.021), but more similar across bands for the <170 Hz narrator (although still some differences, *χ^2^*(4)=17.46, *p*=0.002). More specifically, lowering f0 boosted wave V amplitude by a median of 1.34× larger (IQR: 1.20–1.57×) for the 8 kHz band to 2.42× larger (IQR: 1.98–3.77×) for 0.5 kHz band for the >170 Hz narrator, and by a median of 1.12× larger (IQR: 1.00–1.23×) to 1.26× larger (IQR: 1.07–1.53×) for 8 and 0.5 kHz bands respectively for the <170 Hz narrator. This corresponded to median SNR boosts from the 8 to 0.5 kHz bands of 1.83–6.80 dB for the >170 Hz narrator and 0.58–1.66 dB for the <170 Hz narrator, as well as median speedup ratios of 1.52–4.23× faster for the >170 Hz narrator and 1.14–1.47× faster for the <170 Hz narrator.

**Figure 4.**
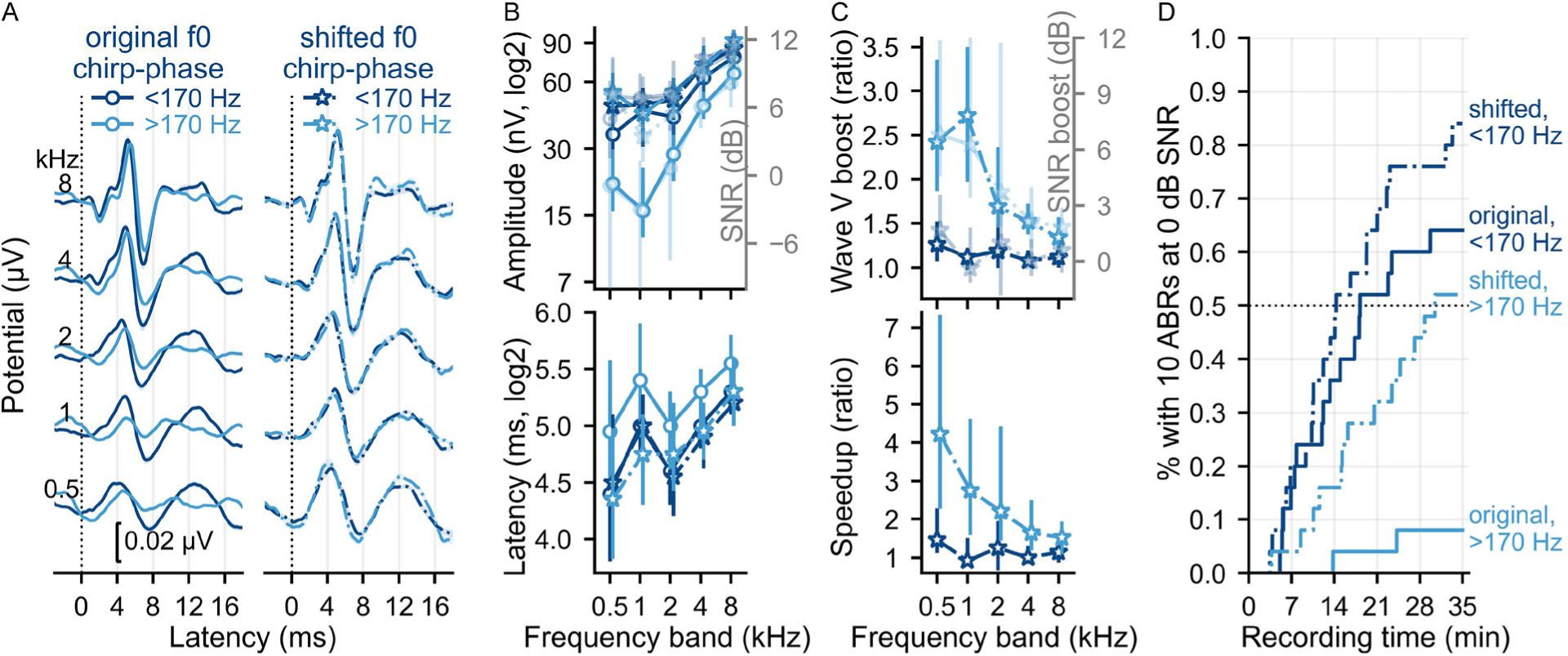
Measured ABR boosts to changing f0 for n=25. (A) Mean ± SE (shading) waveforms for original (solid line) and optimal shifted f0s (dash-dotted line) for stories with original f0s that are lower (darker blue) and higher (lighter blue) than 170 Hz and chirp-phase, along with their corresponding meidan (B) wave V peak amplitudes (top) and latencies (bottom). The signal-to-noise ratio (SNR) over 0-9 ms is also displayed along with the amplitude. (C) Lowered f0 boosts to wave V amplitude and response SNR from 0-9 ms (top) and the associated speedup ratio of testing time calculated from the SNR (bottom). All error bars represent the 25^th^ to 75^th^ percentiles. (D) Cumulative density function of the proportion of participants who had all 10 waveforms (5 bands in 2 ears) reach 0 dB SNR across recording time in minutes. A dotted horizontal line denotes the point at which half the participants achieved this criterion.

These f0-boosts resulted in faster test times for ABRs to reach 0 dB SNR. Lowering the f0 resulted in all 10 waveforms reaching 0 dB SNR a median of 1.97× faster (IQR: 1.00–3.41×) for the >170 Hz narrator and 1.28× faster (IQR: 1.00–1.72×) for the <170 Hz narrator, corresponding to a median of 31.75 minutes saved (IQR: 0–45.2 minutes) for the >170 Hz narrator and 2.00 minutes saved (IQR: 0–21.64 minutes) for the <170 Hz narrator. As shown by the CDFs in Figure 4D, the median test time for the >170 Hz narrator changed from >70 minutes (extrapolated) for the original f0 to 30.47 minutes (IQR: 16.22–57.76 minutes) for the optimal lower f0, and from 18.22 minutes (IQR: 11.97–70.0 minutes) to 14.32 minutes (IQR: 10.29–23.10 minutes) respectively for the <170 Hz narrator. Furthermore, the overall proportion of participants with 10 waveforms at criterion was higher, especially after 20 minutes. Considering the optimal lowered f0 conditions (dot-dashed lines in Figure 4D), the percentage of participants meeting the criterion for the <170:>170 Hz narrators was 64%:28% after 20 minutes and 84%:52% after the maximum of 35 minutes. While less time was saved for the <170 Hz narrator, lowering f0 still conferred benefit by increasing the overall percentage of participants meeting criterion from 64% to 84% after 35 minutes. Thus, when using chirp-phase, the optimal f0 parameter includes a lowered f0 (in this case, 100 Hz) for a narrator with an original f0 <170 Hz.

### Narrators with optimized parameters give similar, but not identical, ABRs and test times

Having verified that multiband peaky speech created with chirp-phase and lower f0s boost ABR responses and cut testing time, we next evaluated whether three different narrators with original f0s of 124, 130 and 147 Hz (i.e., all <170 Hz) can evoke similarly sized ABRs in similar test times when their f0s are lowered to an optimal f0 (90 Hz for this experiment). Figure 5 shows the ABR waveforms (A), wave V peak amplitudes and latencies and response SNR over 9 ms (B), and the CDF for all 10 waveforms to reach 0 dB SNR (C).

**Figure 5.**
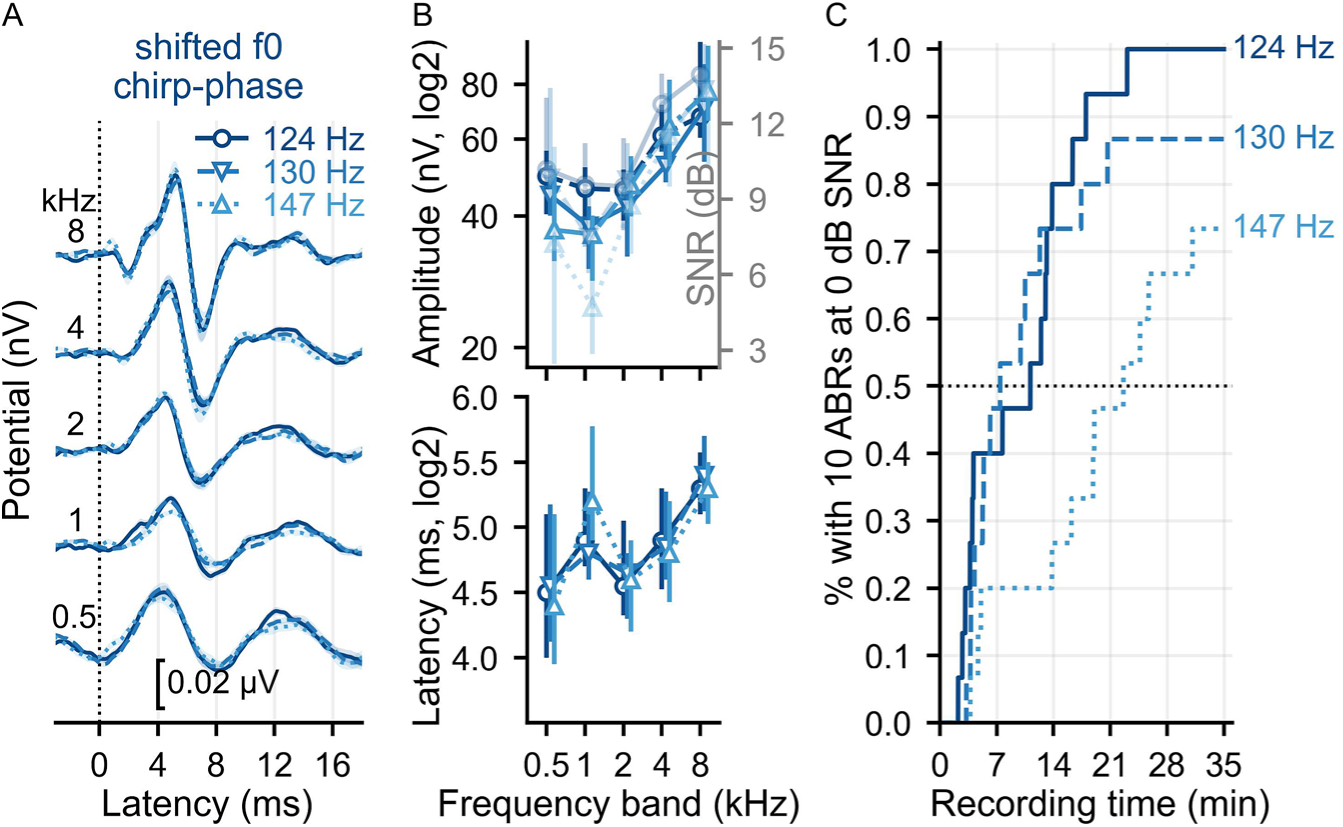
Measured ABRs to 3 audiobooks with chirp-phase and original f0s < 170 Hz that were shifted down to their optimal f0s of 90 Hz for n=15. (A) Mean ± SE (shading) waveforms, along with their corresponding median (B) wave V peak amplitudes (top) and latencies (bottom). The signal-to-noise ratio (SNR) over 0-9 ms is also displayed along with the amplitude. All error bars represent the 25^th^ to 75^th^ percentiles. (C) Cumulative density function of the proportion of participants who had all 10 waveforms (5 bands in 2 ears) reach 0 dB SNR across recording time in minutes. A dotted horizontal line denotes the point at which half the participants achieved this criterion.

With optimized parameters (lowered f0 and chirp-phase), narrators with differing original f0s <170 Hz evoked similar overall ABR waveform morphologies and wave V peak latencies. However, wave V peak amplitudes and response SNRs increased with decreasing original f0 (for all bands: *χ^2^*(2)≤24.27, *p*<0.001; most post-hoc *p*≤0.028). This translated into median test times of 11.15 minutes (IQR: 3.85–13.49 minutes) for the 124 Hz narrator, 7.43 minutes (IQR: 4.76–14.86 minutes) for the 130 Hz narrator, and 22.60 minutes (IQR: 15.01–35.01 minutes) for the 147 Hz narrator (seen as CDF lines crossing the black dotted line in Figure 5C). The narrator with the highest original f0 (147 Hz) had the slowest increase in proportion of participants with all waveforms reaching criterion as recording time increased, whereas the other two narrators had similar proportions of participants over time until ∼15 minutes (73% versus 80%). After 35 minutes, the percentage of participants with all 10 waveforms at 0 dB SNR was 73.3%, 86.7% and 100% for the 147 Hz, 130 Hz, and 124 Hz narrators respectively. While the narrator with the lowest original f0 gave the largest ABRs/SNRs and shortest test times for the most participants, the overall CDFs were similar for the two narrators with the lowest original f0s, suggesting that both could reasonably be used to allow variety in choice of audiobook.

## Discussion

In this study we determined the optimal stimulus and analysis parameters for multiband ABRs to continuous speech in the form of audiobooks. Multiband ABRs are largest and fastest to record to stories with narrators who have original f0s <170 Hz that have peaky speech created with chirp-phase profiles and their f0s shifted down to an optimal range of 90–110 Hz. Modeled ABRs provided good estimates of the measured ABR boosts to parameter changes. The simplest parameter change to chirp-phase cuts the median testing time in half, but also lowering the f0 improves test time and increases the proportion of participants with 10 robust waveforms in under 30 minutes. Different narrators with f0s <170 Hz can be used with optimized parameters, but when possible, the lower the original f0 the more likely participants will have all multiband ABRs in a shorter recording time. Overall, optimizing parameters reduced the testing time from 60 minutes to 14 minutes for ∼70% of participants, making this paradigm more viable as a future assessment tool.

Computational modeling is useful for experimental planning to maximize data quality and efficient testing during the human EEG studies. This approach enabled sampling over a wider f0-phase parameter space to determine the most important conditions to verify in measured responses. The main amplitude trends and boosts were roughly similar between modeled and measured ABRs, suggesting that subcortical processing of the underlying acoustics accounted for a large portion of parameter effects on the ABRs. Furthermore, the modeled ABRs helped identify which of the 12 selected narrated stories would evoke large ABRs, as well as which lowered f0 was optimal for each story (Supplemental Figure 1). This in turn helped build a set of potentially effective stories for use in experiments. The slightly lower boosts for the measured ABRs could reflect differing recording conditions (e.g., noise) and individual factors such as hearing thresholds and residual integrity of the cochlear components contributing to neural synchrony. We only screened participants for normal hearing at 20 dB HL and used a flat 0 dB HL audiogram for the modeled ABRs. Next, we could measure actual hearing thresholds and model different audiograms to further investigate the extent to which variability in normal hearing contributes to the range of expected ABR outcomes. Although the overall pattern was properly captured, the pattern of ABR wave V latencies slightly differed across octave bands. Consistent with this, other studies have reported that latency predictions are less accurate than amplitude predictions, and suggested that complex stimuli may not be as accurately represented by the constant brainstem delays assumed by the models (Dau, 2003; Temboury-Gutierrez et al., 2024). The models also do not account for cortical activity, although our analysis step of subtracting the common component should limit the cortical contributions to our multiband ABRs, as would our higher 80 Hz high-pass filter cutoff. Despite these minor differences between modeled and measured responses, the overall patterns provided good enough estimates to be helpful for teasing apart some acoustic-based effects of changing parameters and checking complex speech stimuli before conducting an experiment.

Changing both phase and f0 optimizes ABRs to multiband peaky speech. However, just changing the phase profile alone from zero- to chirp-phase evoked substantially larger ABRs and faster testing times, bringing the necessary testing times for 70% of participants down from 60 minutes to 20 minutes (Figure 3D) and the median testing time to ∼10 minutes. This parametric change is easy to implement and does not affect the spectrotemporal dynamics, resulting in little, if any, perceptual change to the peaky speech stimuli. Therefore, if it’s important for the scientific question to keep f0 and formant relationships unaltered (beyond the small random f0 shifts necessary for creating multiband speech), chirp-phase is a viable single parameter change for evoking higher quality ABRs in less time. Accounting for the cochlear delays improved neural synchrony especially at the cochlear apex, evidenced by the greatest amplitude increases to the lowest 0.5 kHz octave band (Figure 3B,C). The 0.5 kHz octave band was the broadest and smallest response and primary contributor to delayed test times for the zero-phase stimuli (Polonenko & Maddox, 2021). Unsurprisingly, the chirp-phase wave V latencies were more similar across octave bands than for zero-phase stimuli, reflecting compensation for cochlear delays. But interestingly, the wave V latencies still differed by band (Figure 3B, bottom row), and in fact became shorter for lower- than higher-frequency bands, suggesting that the CE-chirp may have over-estimated the cochlear delays for speech stimuli. The CE-chirp was created based on derived-band ABRs (Elberling et al., 2007) and we used the level-independent version as a starting point because the correspondence between levels of narrowband chirps and our speech bands was not obvious. Other chirps, such as the level-dependent CE-chirp (Elberling & Don, 2010) or new chirps created based on speech stimuli, may better account for the cochlear delays to continuous speech and consequently evoke even larger ABRs. Although, any additional amplitude boosts and time saved from different chirps are likely to be subtle compared to the main zero- versus chirp-phase boosts seen here.

Likewise, lowering the f0 evokes larger responses and improves testing time, but also allows for the use of a wider range of narrators. The ABR boosts for lower f0s is consistent with previous work on ABRs to broadband peaky speech and click stimuli with stimulation rates matching the speech f0s (Polonenko & Maddox, 2024). Faster glottal pulse rates (i.e., higher f0) behave similar to higher click stimulation rates (Polonenko & Maddox, 2024), which evoke smaller responses due to neural refractoriness and adaptation (Burkard et al., 1990; Burkard & Hecox, 1983; Chiappa et al., 1979; Don et al., 1977; Jiang et al., 2009). Thus, lower f0s evoke greater neural synchronization and larger ABRs, which in turn allows faster testing time. Lowering f0 saved relatively more time for the narrator with an original f0 >170 Hz, bringing the ABR responses and testing times into the range of the other narrator’s original f0 responses, thereby making it a more effective stimulus. The boosts to ABR amplitudes and SNR were more modest and similar across bands for the narrator with an original f0 <170 Hz compared >170 Hz (Figure 4C), but were still effective at decreasing test time (Figure 4D). This was particularly evident for the “slower” half of participants that required longer test times with the original f0 stimuli (where the dark blue CDFs diverge in Figure 4D). Unlike the ∼1.3× speedup and ∼2 minutes saved for the “faster” half of participants, lowering f0 saved at least 21.6 minutes (at least 1.7× faster, 75^th^ percentile). These gains from lowering f0 meant that 75% of participants went from requiring 35 minutes to 21 minutes for the <170 Hz narrator. Furthermore, these improved test times were not just limited to the specific story: other stories with optimized parameters evoked similarly sized ABRs and required ∼21-minute test times for most participants (Figure 5C). Some differences remained between the various narrated stories, which could be due to the small differences in original f0s, as narrators with progressively lower original f0s had the best SNRs and test times (Figure 5B,C). But other factors may contribute as well, such as greater f0 variation from imitating different characters in the story or differing overall spectral power in each octave band. Future studies could investigate additional narrator parameters to further maximize multiband testing and widen story selection options to allow a corpus of stimuli that may be engaging across different participant preferences.

Analysis parameters are also important for optimizing the multiband peaky speech paradigm. We systematically evaluated different filter parameters and SNR windows to maximize response metrics. The high-pass cutoff particularly affects response amplitude and morphology, and thus, the appropriate cutoff will depend on which components of the response are the focus of evaluation. Our previous work with peaky speech suggests a 150 Hz cutoff allows better visualization of earlier waves I and III, whereas later MLR waves are better viewed with a 30 Hz cutoff (Polonenko & Maddox, 2021). For the purpose of determining the presence of a response in each octave band, we focused on the largest wave V component. Using data from 3 experiments, we determined that an 80 Hz cutoff gave the largest responses and SNRs across conditions and participants. An 80 Hz cutoff may also generalize better to pediatric populations than higher cutoffs (e.g., 150 Hz) because their ABRs have a greater proportion of lower energy than adult ABRs (Spivak, 1993), which is why lower high-pass cutoffs are often used in frequency-specific diagnostic ABR exams in infants (American Academy of Audiology, 2020; Hatton et al., 2022; Ontario Ministry of Children and Youth Services, 2020).

We also evaluated the optimal SNR analysis window. Previously, a 0–15 ms window was used that covered the entire ABR response and component waves (Polonenko & Maddox, 2021). To fairly represent the zero-phase latency differences between octave bands while mainly covering parts of the response that vary most (i.e., wave V peak and trough), we shifted the starting latencies of the SNR windows for the zero-phase responses, as done with other frequency-specific tone pip ABR studies (Polonenko & Maddox, 2019, 2022). A 9 ms window was optimal across parameters and participants, although 8.5–9.5 ms gave reasonably similar overall SNRs and test times. Of particular importance, the response SNRs calculated over the 9 ms window followed the same patterns across octave bands as wave V peak amplitudes (Figure 3B, Figure 4B, Figure 5B). For studies focusing on wave V or the presence of a response, SNR may provide a better metric than peak amplitude because it gives the same trends while accounting for background noise and avoiding any pitfalls of automatic or manual peak-picking.

There are other ways that may further speed up testing and allow more variety in audiobooks. As mentioned above, other chirp profiles may better compensate for cochlear delays to complex speech stimuli. Ideally, all 10 audiological bands will be measured, but our previous work shows that 8 bands reach criterion SNR in less time (Polonenko & Maddox, 2021). To reduce the number of bands, we could combine the two lowest or highest octaves or remove the 8 kHz bands―which are currently not tested in diagnostic ABRs―by lowering the upper filter cutoff for the re-synthesized speech. Often suprathreshold ABR tests are conducted at higher levels, especially when earlier component waves are analyzed. We could also use a higher level than the ∼65 dB SPL conversational level used here, which might be more tolerable for shorter test times than over the ∼2 hours required for these experiments.

In summary, optimized stimulus and analysis parameters of the multiband peaky speech paradigm improve ABR response sizes to allow more feasible testing times. Chirp-phase boosts the most difficult to measure low-frequency octave band responses without changing the spectrotemporal dynamics of the speech. Lowering f0 to 90–110 Hz increases the variety of narrated audiobooks that effectively evoke larger ABRs, thereby allowing a wider variety of options to engage individuals. Combined together, these parametric changes further developed the multiband peaky speech paradigm into a more efficient assessment tool.

## Acknowledgments

We thank Samantha Krocak, Natalie Falconer, and Jenna Skare for their assistance with data collection. We also thank Liberal Arts Technologies and Innovation Services at the University of Minnesota for use of the high-performance computing cluster.

## Statements and Declarations

### Ethical considerations

This study was approved by the Institutional Review Board of the University of Minnesota (STUDY 17008) on September 22, 2022. This research was conducted ethically in accordance with the World Medical Association Declaration of Helsinki.

### Consent to participate

All participants provided written informed consent prior to enrolment in the study.

### Consent for publication

Informed consent for publication was provided by the participants.

### Author Contributions

MP – Conceptualization, Formal analysis, Funding acquisition, Investigation, Methodology, Supervision, Visualization, Writing – original draft, Writing – review & editing

BE – Methodology, Resources, Software, Visualization, Writing – review & editing

