## Supplemental Figure for "Optimal parameters for measuring multiband auditory brainstem responses to continuous speech"

### Optimal parameters for speech multiband ABRs: Supplemental Figures

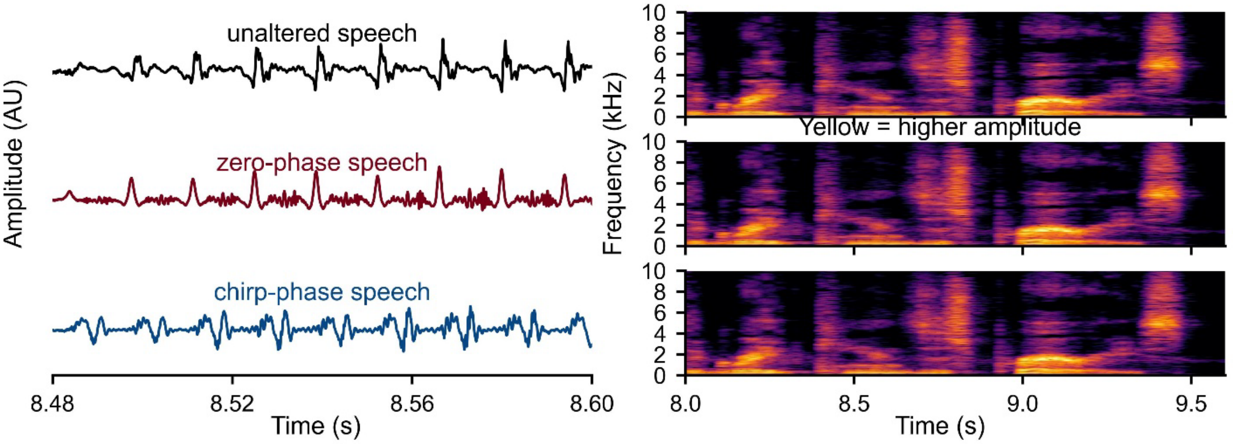


*Supplemental Figure 1. Stimulus examples. The waveforms (left) for the left ear shows that zero-phase multiband peaky speech is more "click-like" than original speech and the high-frequencies start before the low-frequencies for the chirp-phase speech, while their spectrograms (right) are nearly identical. Light yellow colors represent larger amplitudes*

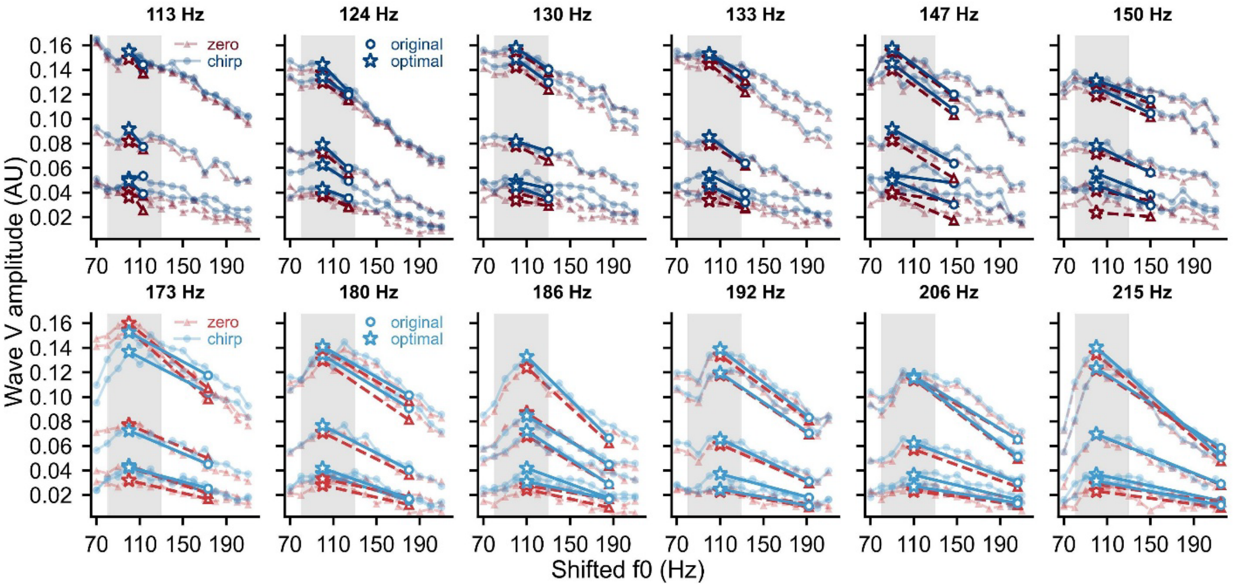


*Supplemental Figure 2. Modeled ABR wave V amplitudes for speech stimuli with the f0 shifted from 70 to 210 Hz in 10 Hz steps. The optimal range is in shaded gray, and the larger shapes denote the original f0 and optimal shifted f0 for each story and phase. Each line represents a frequency band, with lower frequency bands at the bottom and higher frequency bands at the top of each panel.*

### Automatic peak-picks

#### Modeled ABRs

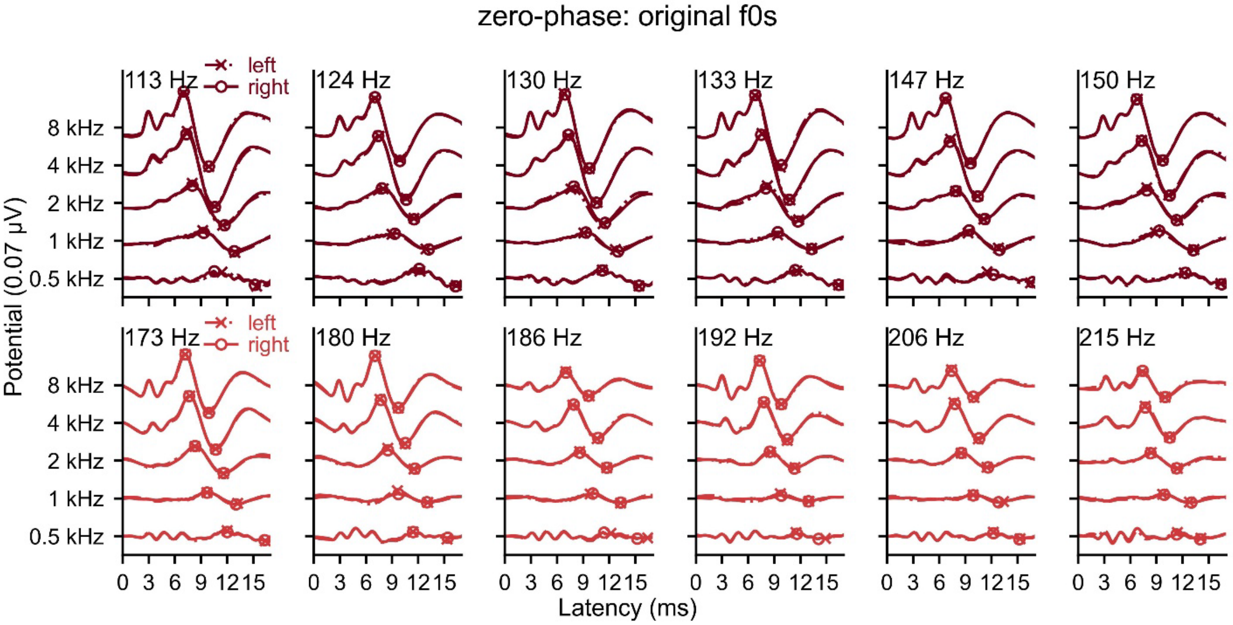


*Supplemental Figure 3. Modeled ABRs and automatic wave V picks for speech stimuli with original f0s and zero-phase.*

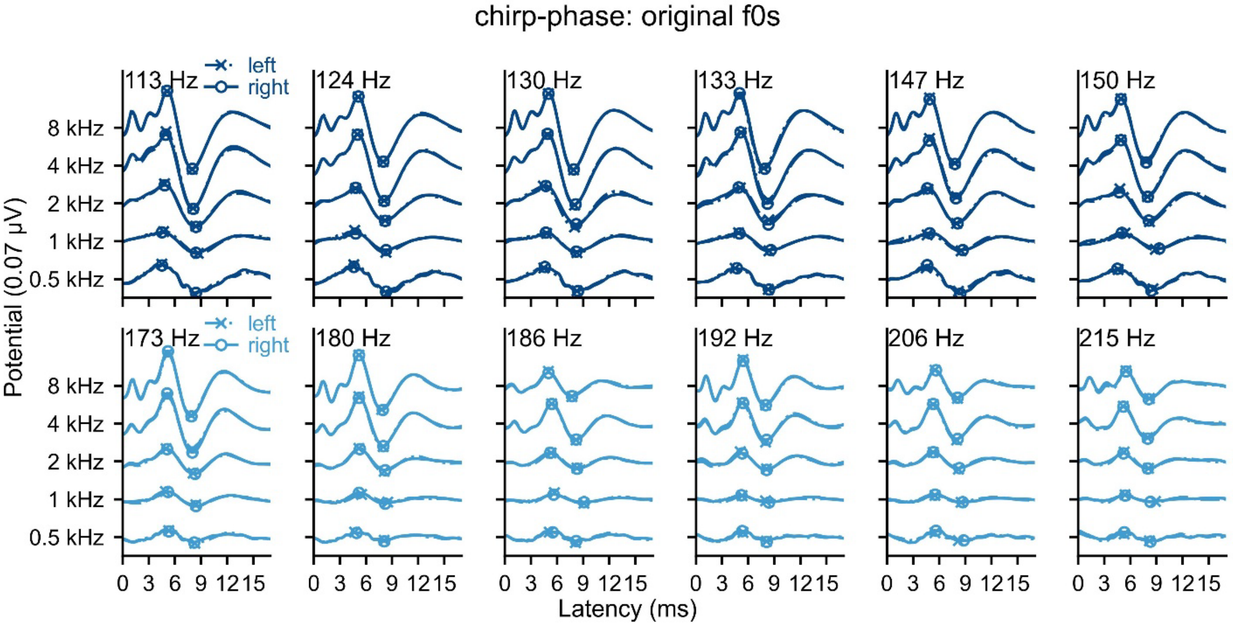


*Supplemental Figure 4. Modeled ABRs and automatic wave V picks for speech stimuli with original f0s and chirp-phase.*

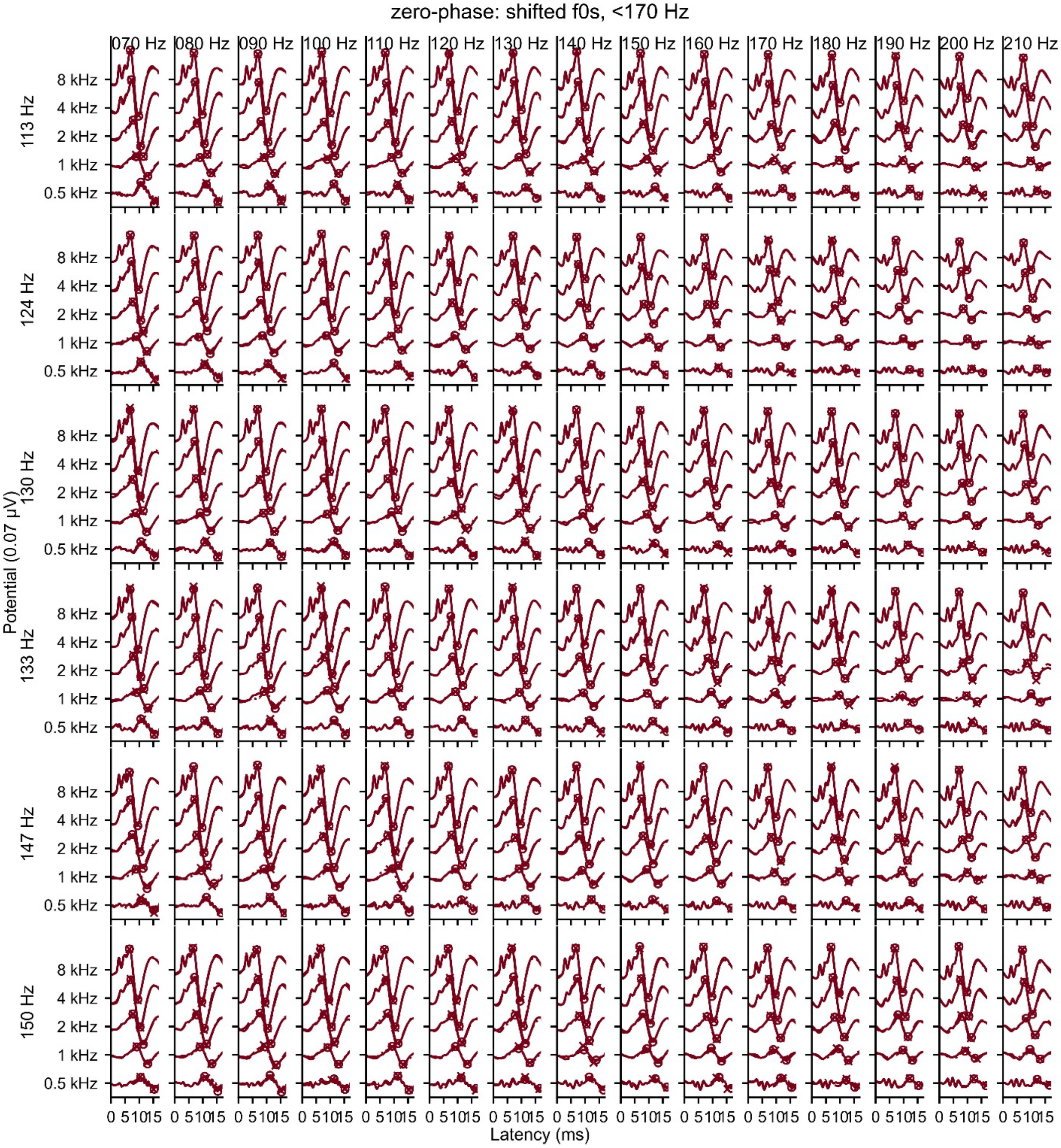


*Supplemental Figure 5. Modeled ABRs and automatic wave V picks to shifted f0s for audiobooks with original f0s <170 Hz and zero-phase.*

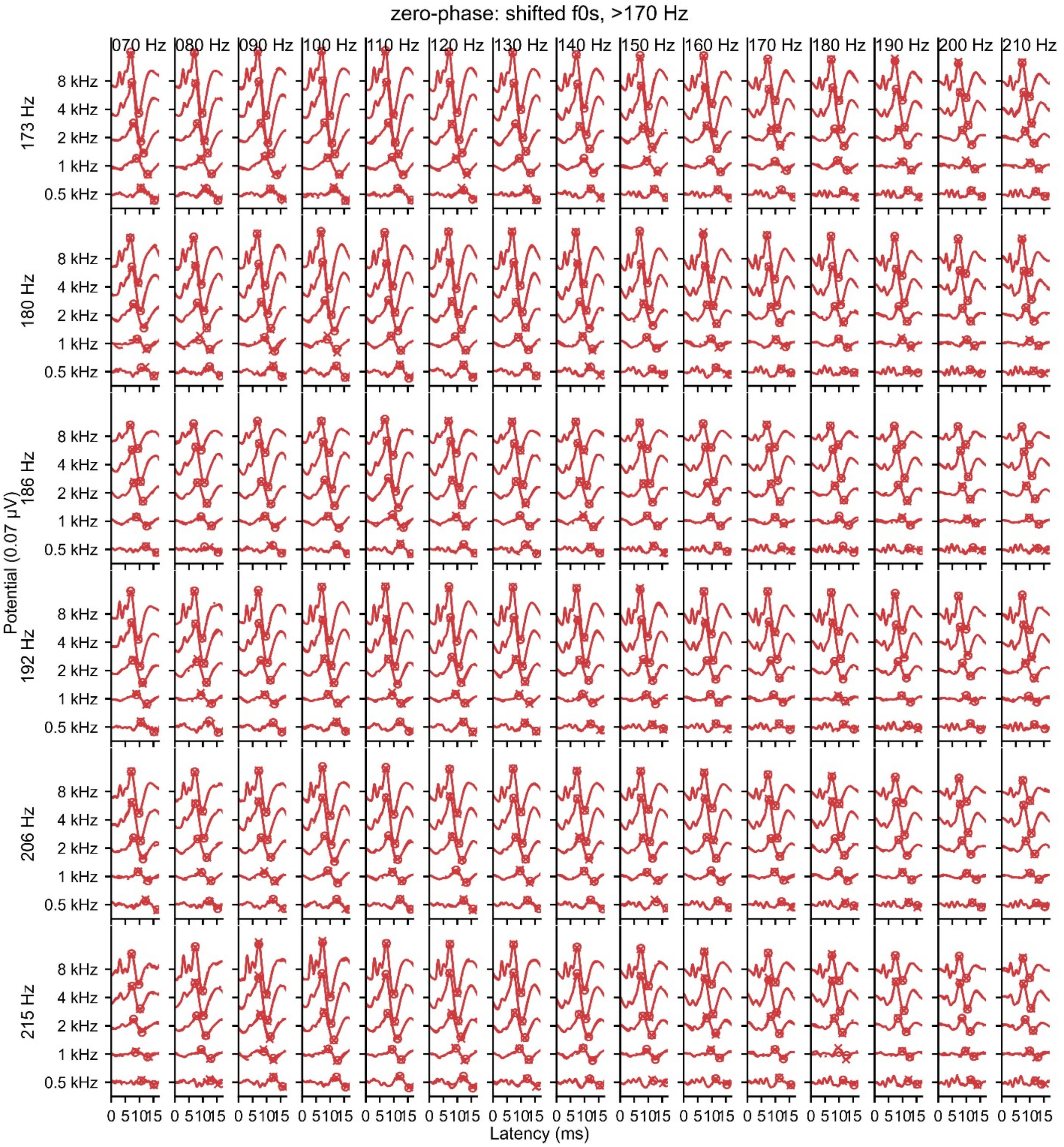


*Supplemental Figure 6. Modeled ABRs and automatic wave V picks to shifted f0s for audiobooks with original f0s >170 Hz and zero-phase.*

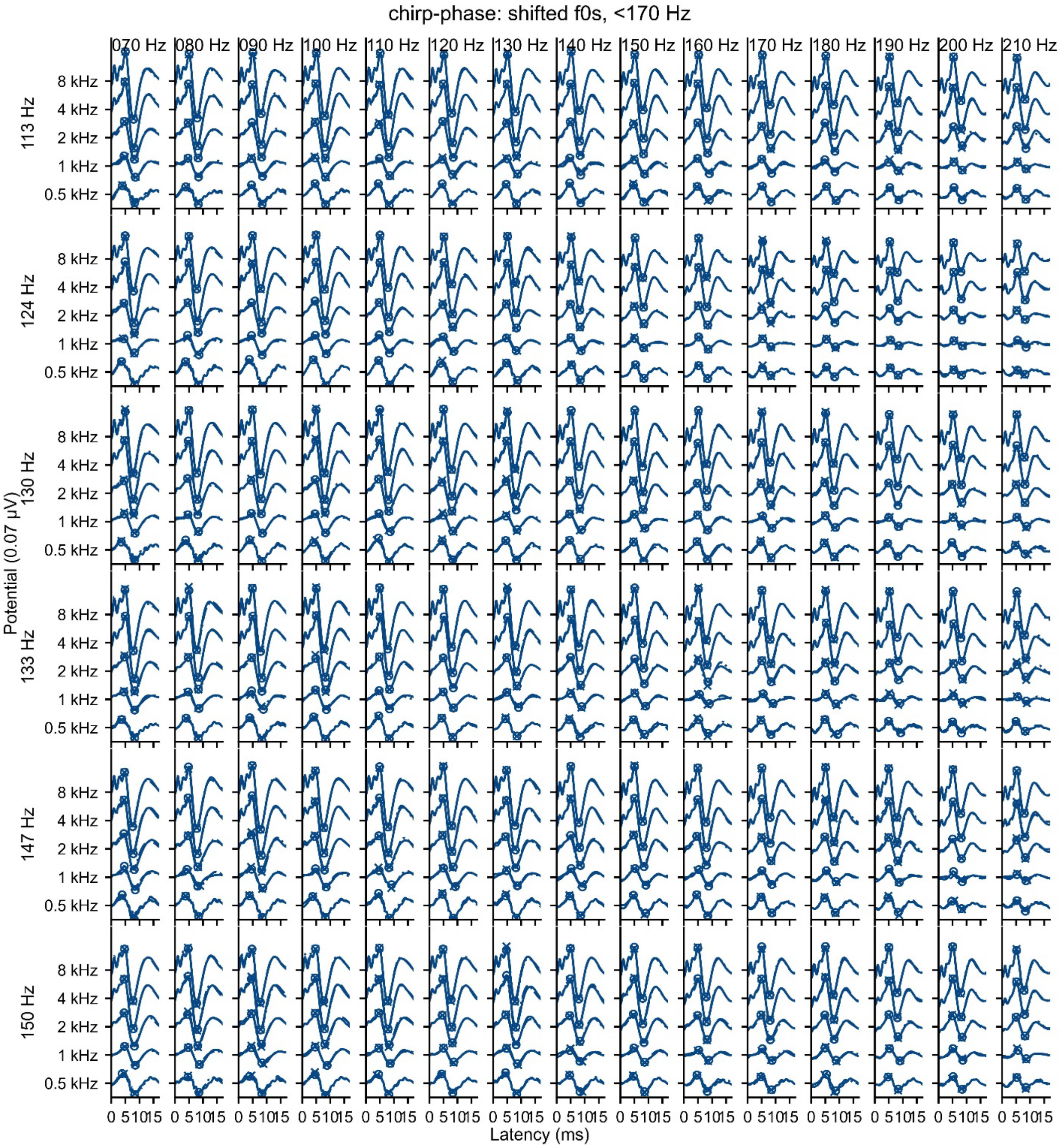


*Supplemental Figure 7. Modeled ABRs and automatic wave V picks to shifted f0s for audiobooks with original f0s <170 Hz and chirp-phase.*

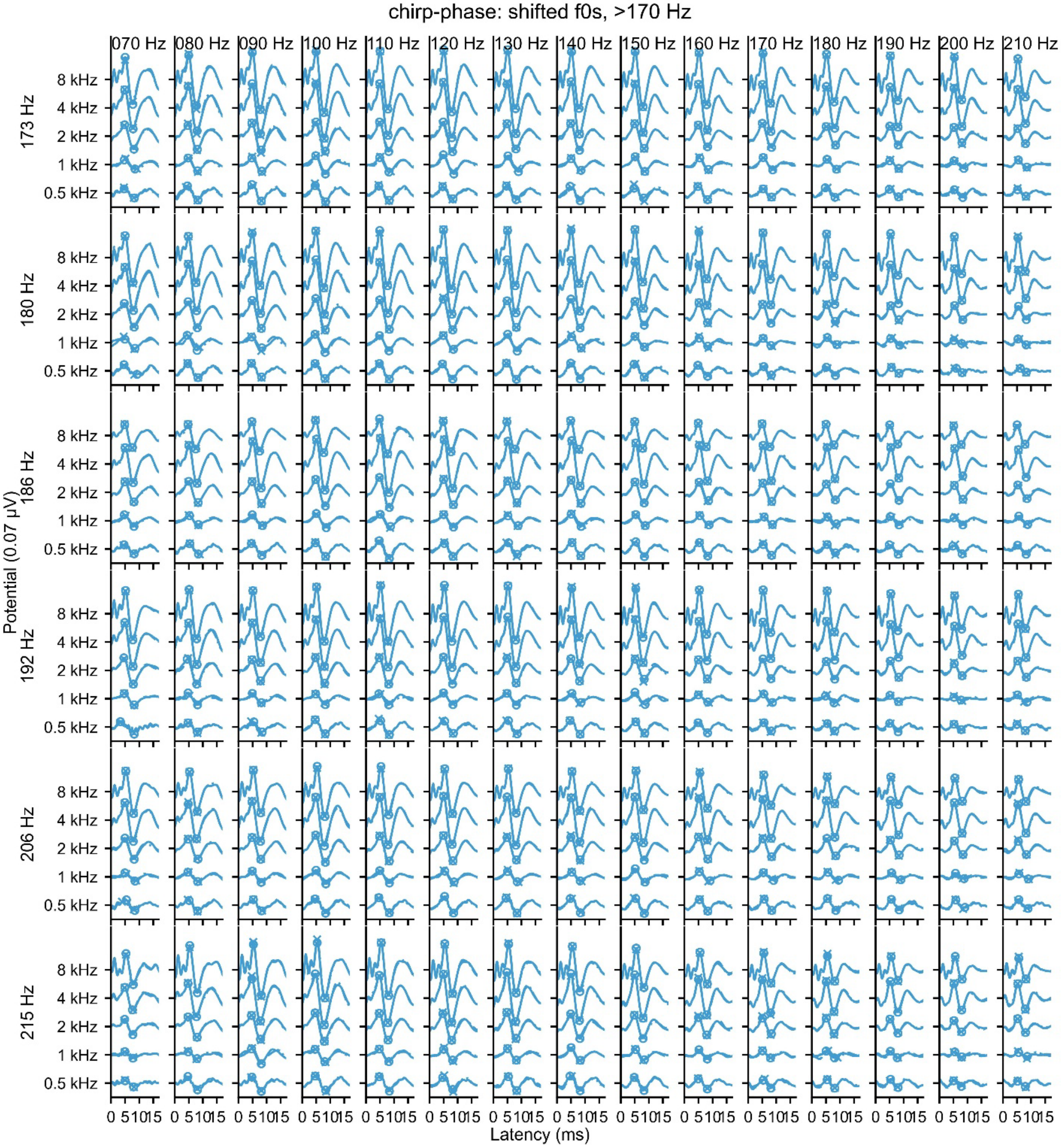


*Supplemental Figure 8. Modeled ABRs and automatic wave V picks to shifted f0s for audiobooks with original f0s >170 Hz and chirp-phase.*

#### Measured ABRs

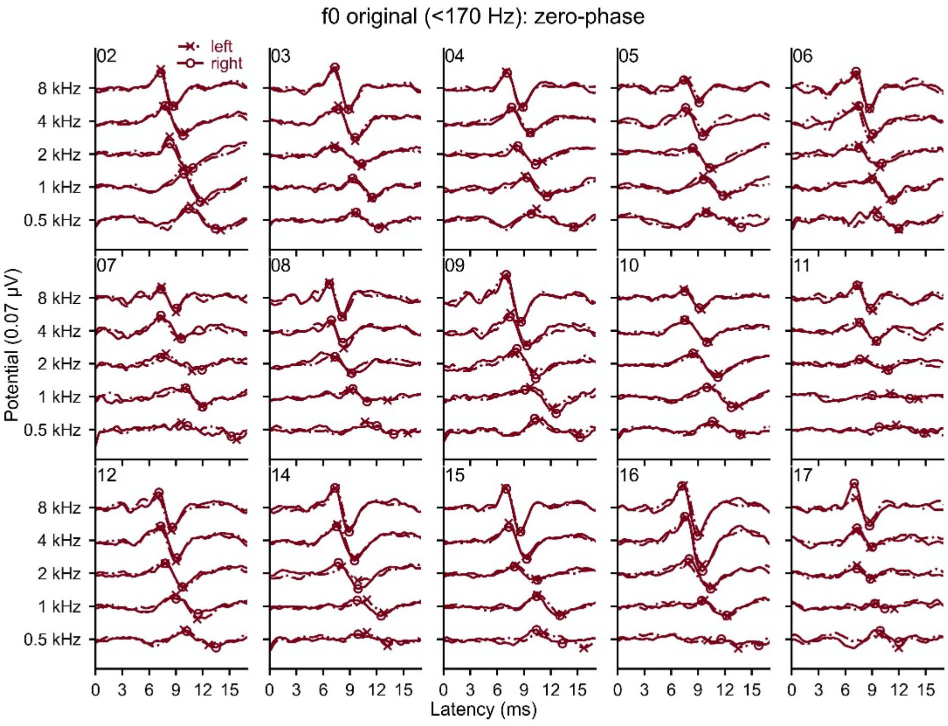


*Supplemental Figure 9. Measured ABRs and automatic wave V picks for zero-phase for each participant.*

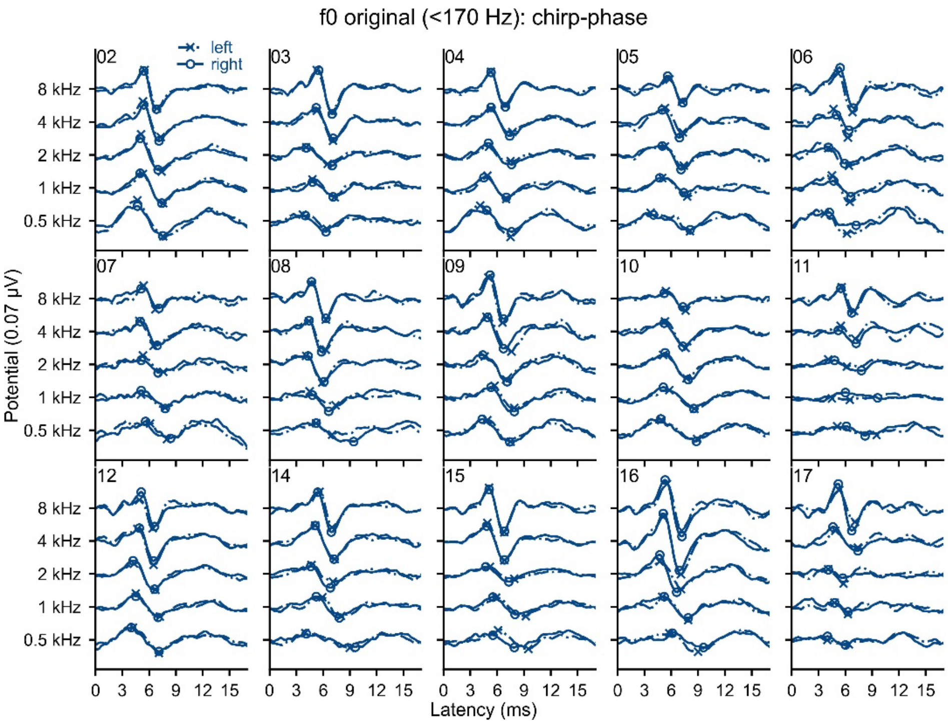


*Supplemental Figure 10. Measured ABRs and automatic wave V picks for chirp-phase for each participant.*

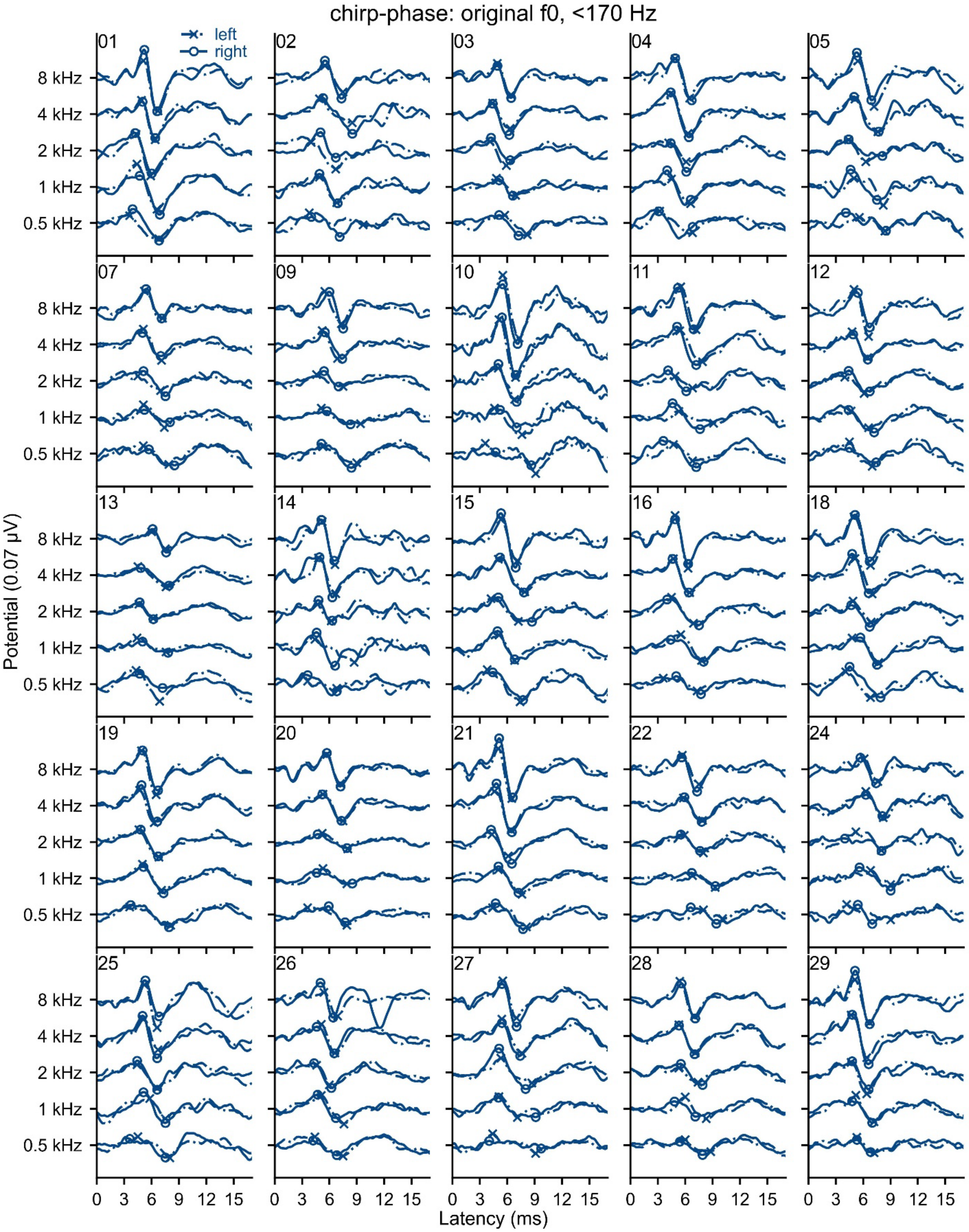


*Supplemental Figure 11. Measured ABRs and automatic wave V picks for each participant to speech with chirp-phase and an original f0 <170 Hz.*

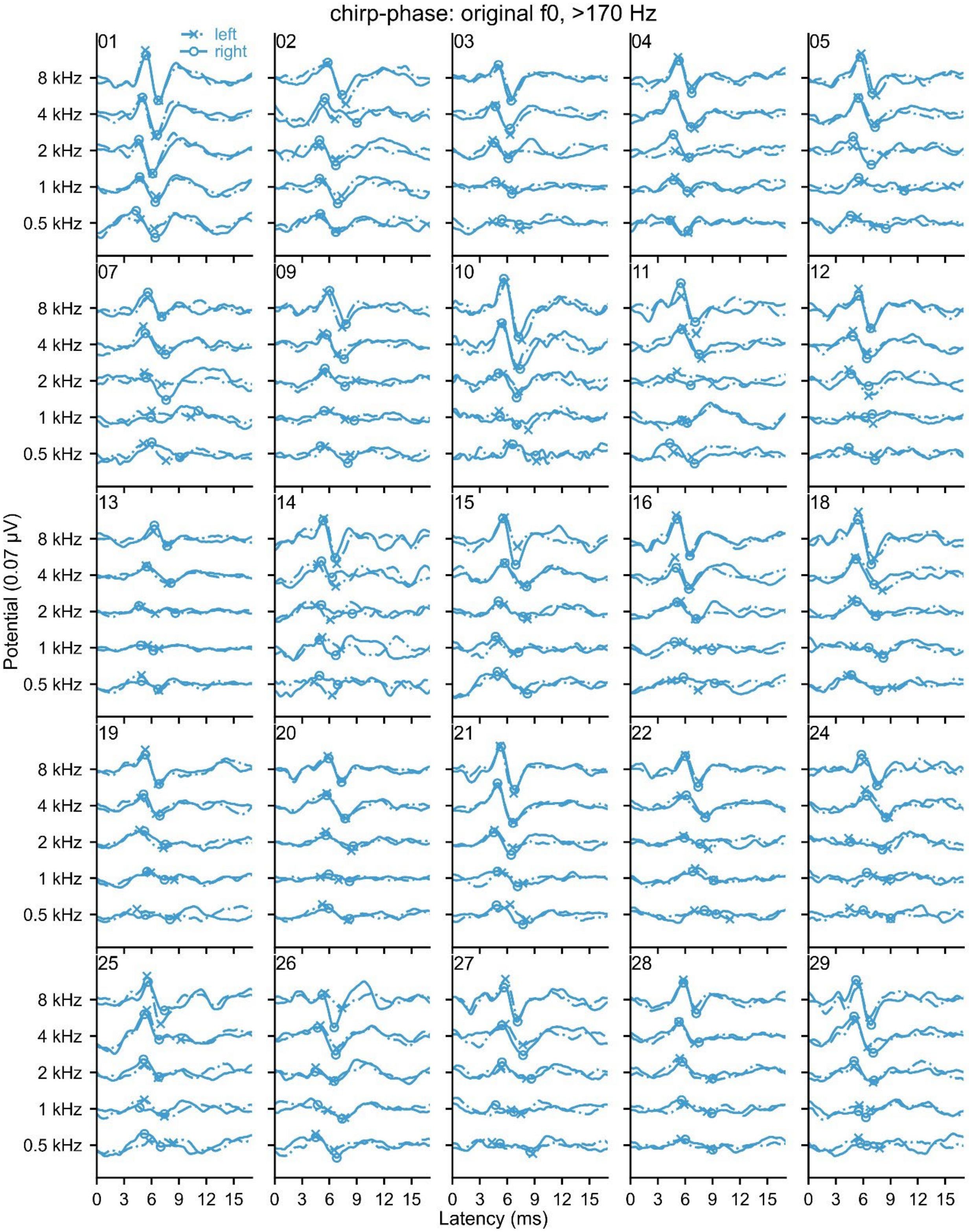


*Supplemental Figure 12. Measured ABRs and automatic wave V picks for each participant to speech with chirp-phase and an original f0 >170 Hz.*

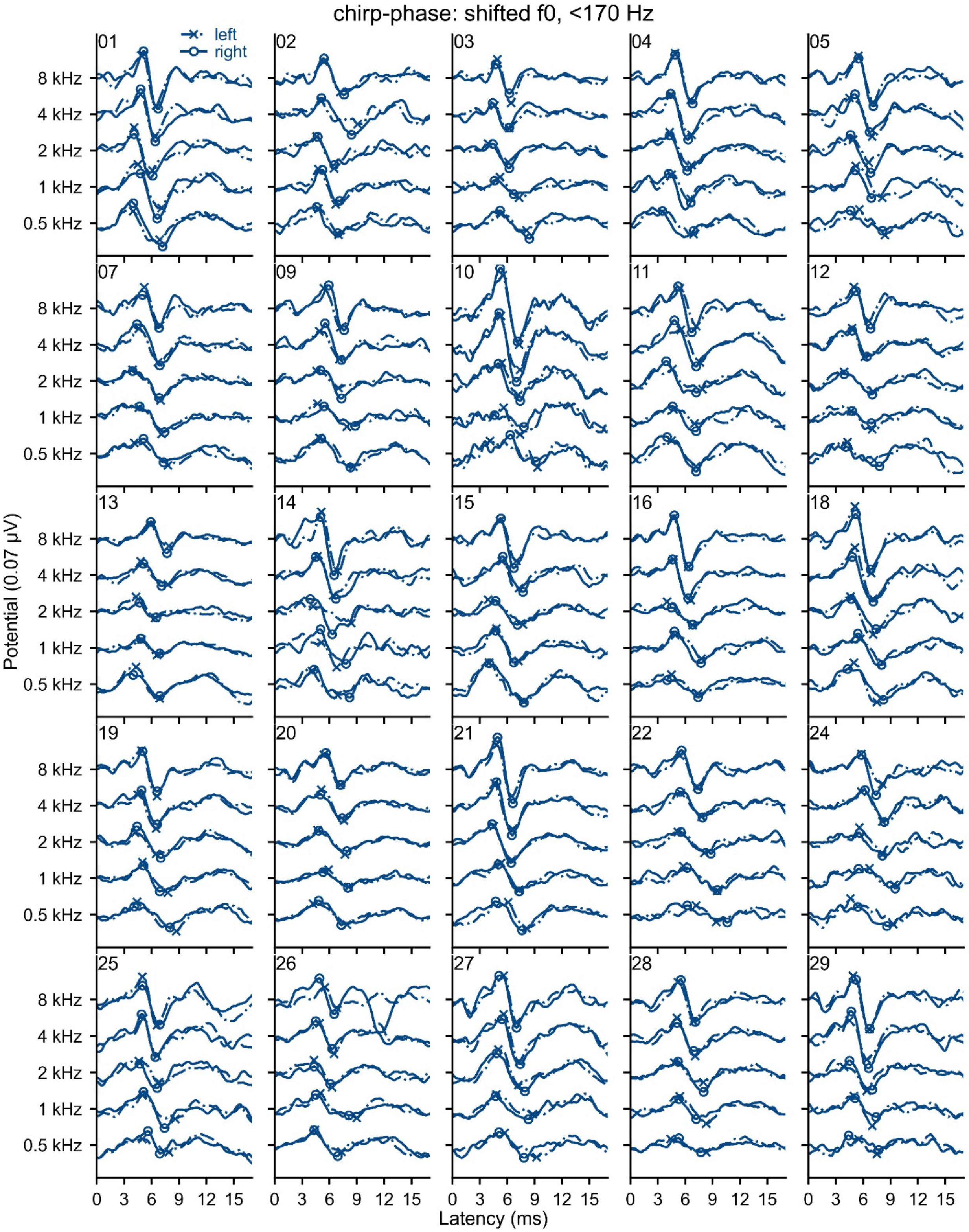


*Supplemental Figure 13. Measured ABRs and automatic wave V picks for each participant to speech with chirp-phase and an original f0 <170 Hz that was shifted down to 90 Hz.*

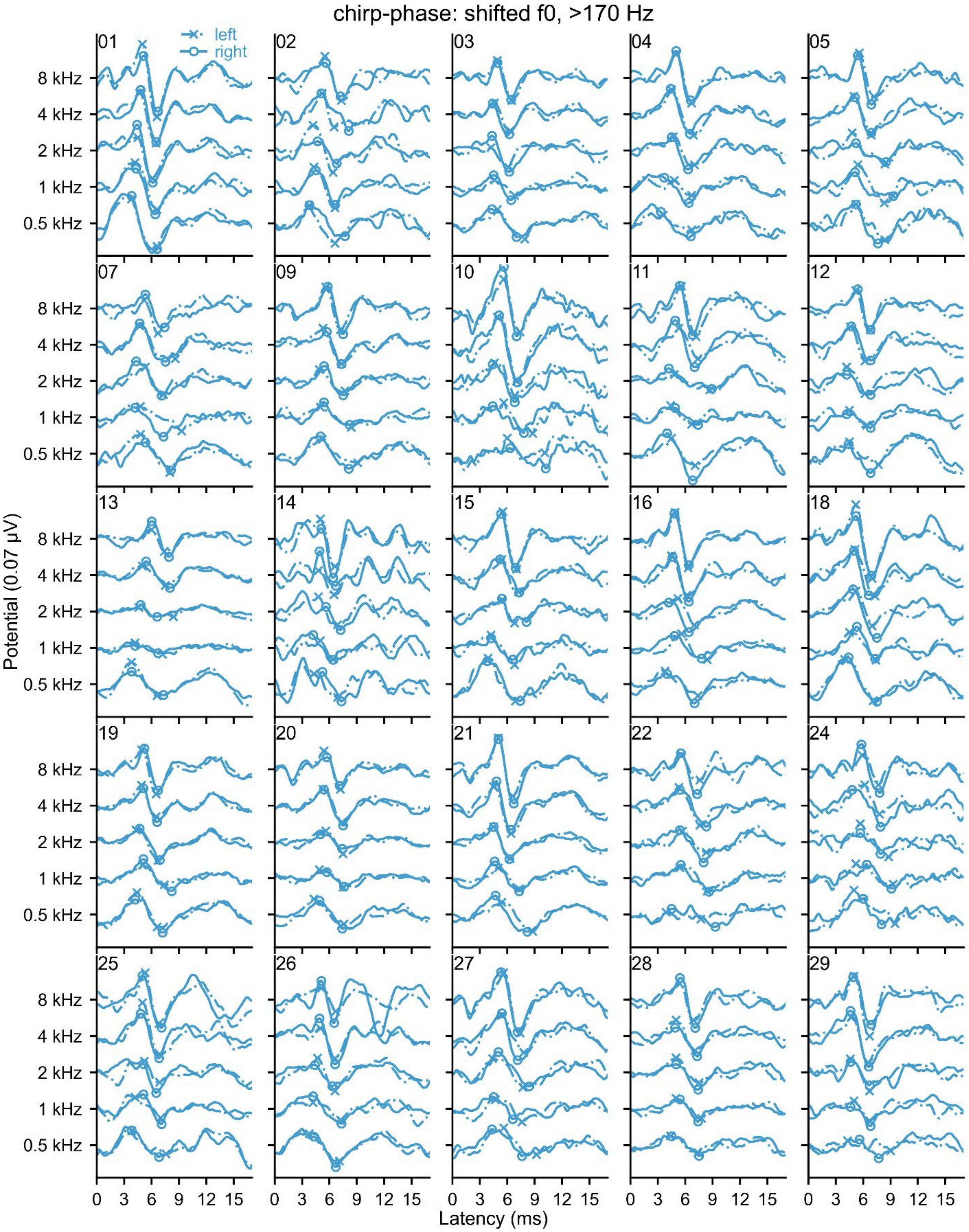


*Supplemental Figure 14. Measured ABRs and automatic wave V picks for each participant to speech with chirp-phase and an original f0 >170 Hz that was shifted down to 90 Hz.*

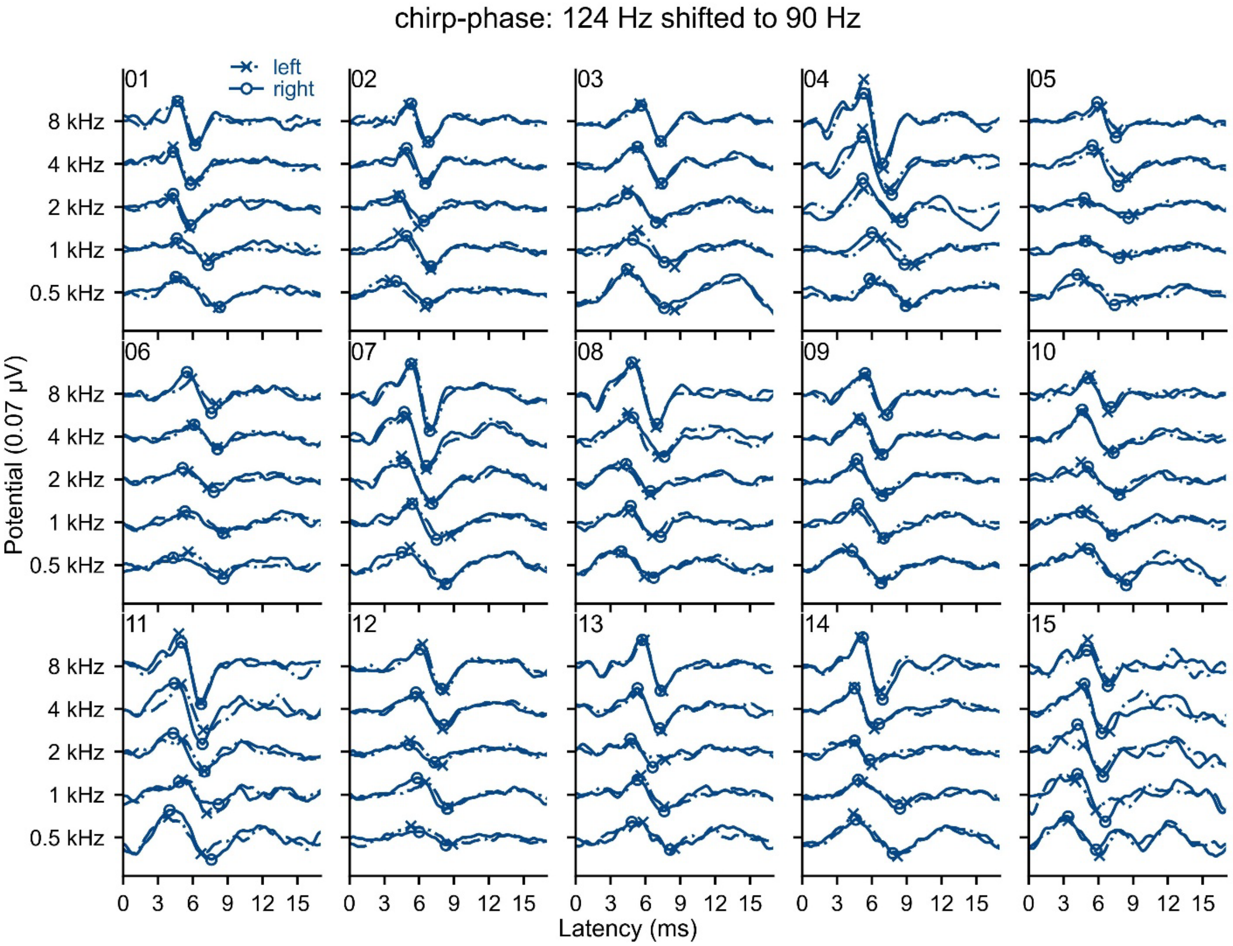


*Supplemental Figure 15. Measured ABRs and automatic wave V picks for each participant to speech with chirp-phase and an original f0 of 124 Hz that was shifted down to 90 Hz.*

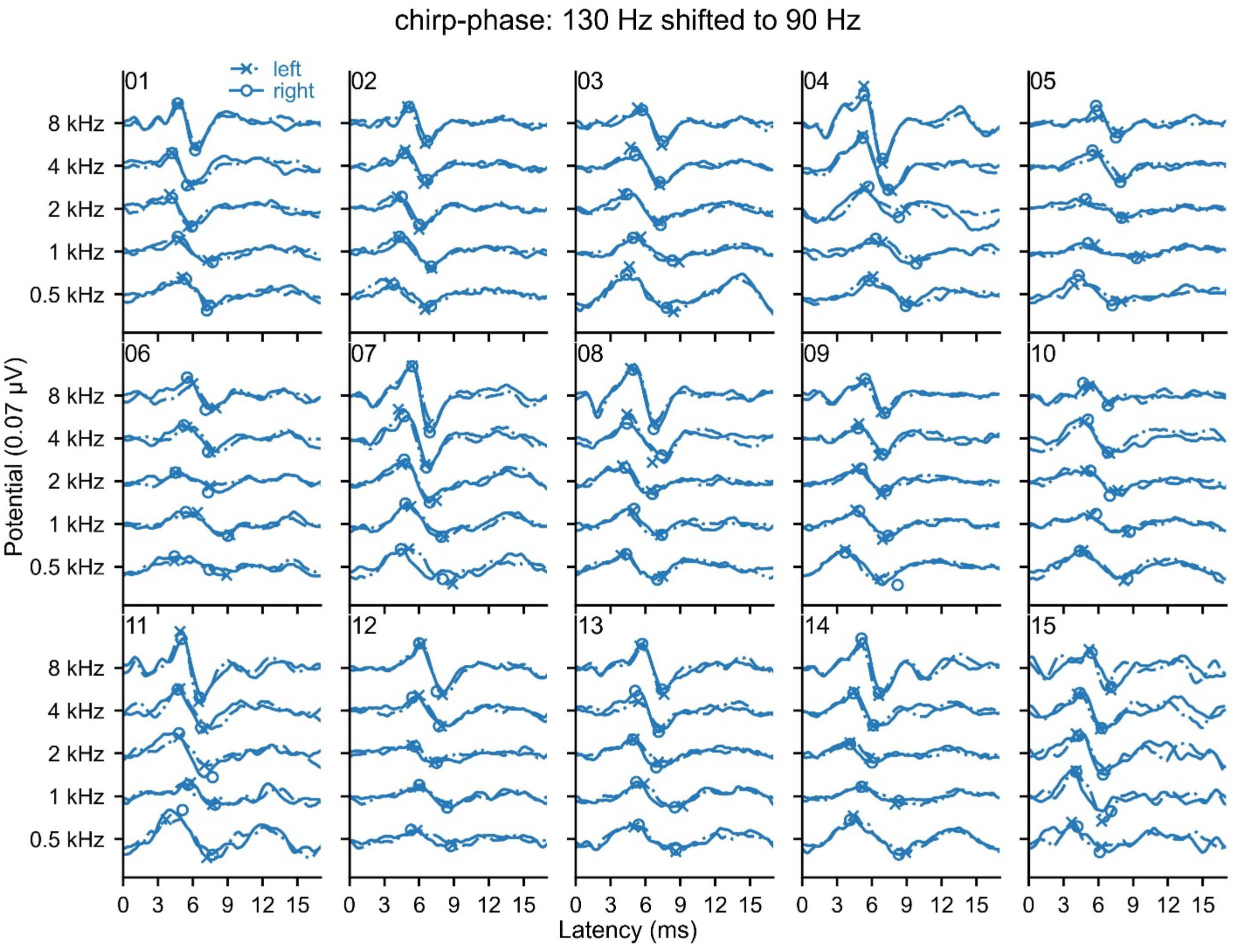


*Supplemental Figure 16. Measured ABRs and automatic wave V picks for each participant to speech with chirp-phase and an original f0 of 130 Hz that was shifted down to 90 Hz.*

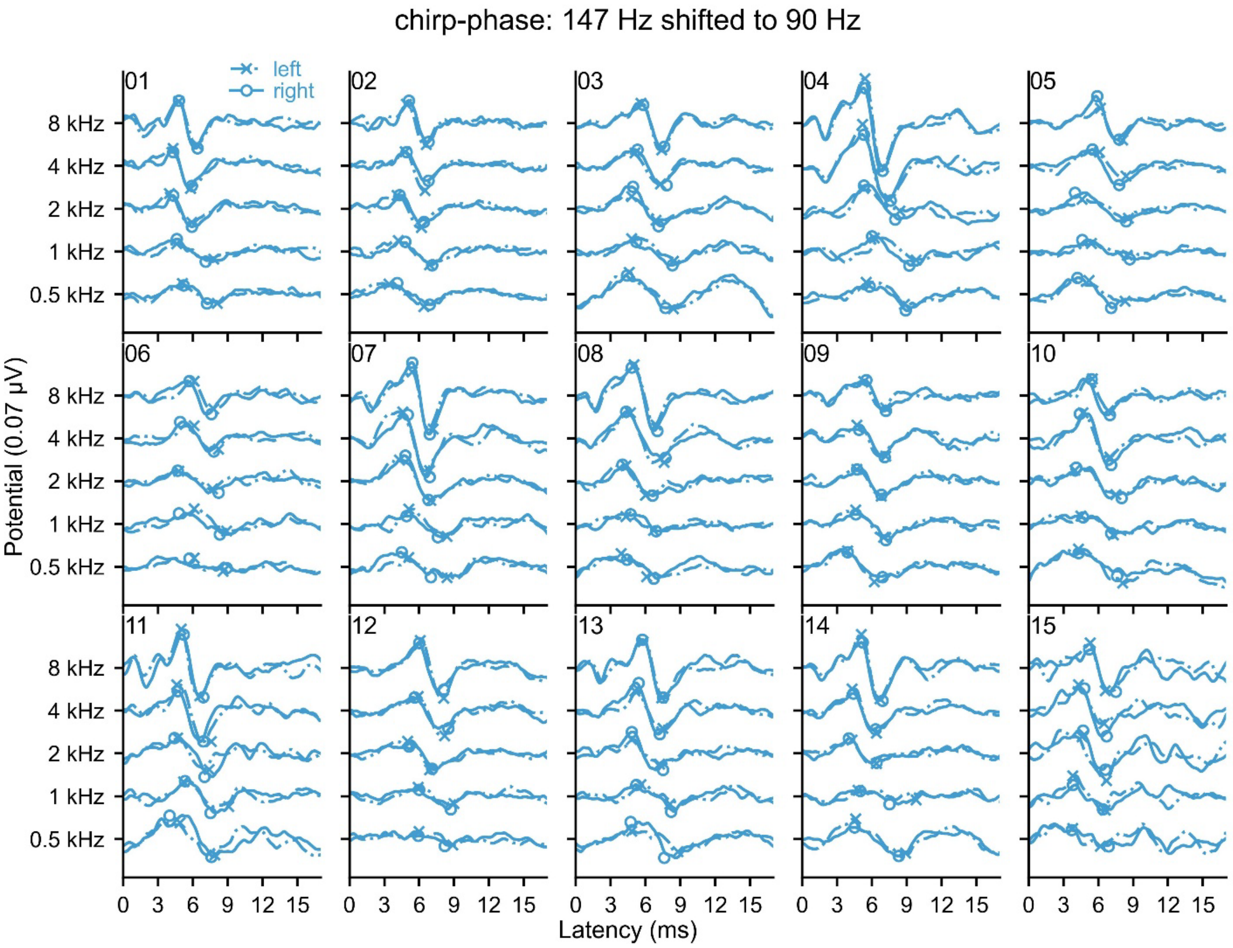


*Supplemental Figure 17. Measured ABRs and automatic wave V picks for each participant to speech with chirp-phase and an original f0 of 147 Hz that was shifted down to 90 Hz.*
